# Morphomechanic tuning of ERK by actin-TFII-IΔ regulates cell identity

**DOI:** 10.1101/2023.06.02.543427

**Authors:** Qiao Wu, Jian Zhang, Bing Long, Xiao Hu, Bruna Mafra de Faria, Stephen Maxwell Scalf, Kutay Karatepe, Wenxiang Cao, Nikolaos Tsopoulidis, Andres Binkercosen, Masaki Yagi, Aaron Weiner, Mary Kaileh, Enrique M. De La Cruz, Ananda L Roy, Konrad Hochedlinger, Shangqin Guo

## Abstract

Cell morphology is faithfully coupled to its identity but the coupling mechanism remains elusive. Using somatic cell reprogramming into pluripotency as a model system, we show that activity of the extracellular signal-regulated kinase (ERK) is tuned by cellular morphomechanic state to direct cell fate. Pluripotent cells and somatic cells reprogramming into pluripotency allocate large amounts of actin into their nucleus, which morphs cells to become taller than 10 μm, a minimal height required for the pluripotent identity. Accumulated nuclear actin binds to TFII-IΔ, an atypical transcription factor that translocates into the nucleus upon signaling. TFII-IΔ also binds to and activates ERK. The binding of TFII-IΔ by nuclear actin reduces ERK activity, in coordination with changes in cell/colony height. The tight coupling between cell height and nuclear actin accumulation necessitates the degree of ERK tuning to be mild. Mild ERK inhibition by chemicals recapitulates the tuning by actin-TFII-IΔ and turns most cells in reprogramming cultures into pluripotency. Thus, we uncover a novel mechanism for how cell morphology couples to its identity via the actin-TFII-IΔ-ERK axis, identifying points of intervention in cell fate manipulation.

## Introduction

Cell morphology is faithfully coupled to its identity, as abnormalities in the former form the basis of histopathology. While the cell form-identity relationship holds true, how these two parameters are mechanistically coupled remains unclear. Furthermore, specific cell morphology is often seen as the product of gene expression programs, passively serving as an indicator of cell identity. Given the fidelity of this form-identity couple, however, it is possible that cell morphology plays a more active role in setting cell identity. Insights into these questions could help the rational design of cell fate engineering approaches and developing more effective intervention strategies, as morphomechanic cues could be implemented in culture protocols or by engineering approaches and devices ^1,2^.

Cell morphology has been mostly examined in the x-y dimension, e.g. by patterning the size/shape of the surface area on which cells adhere and grow ^1,3,4^, or by physically stretching the surface that cells grow on ^5,6^. The significance of cell’s z dimension has only begun to be appreciated when cell height confining devices were implemented, although these efforts focused on short time scales (minutes) insufficient for most cell fate changes ^7,8^. Recent years have also witnessed the explosion of 3D culture techniques (e.g. spheroids, organoids, embryoids and more) that capture cell fates far beyond those attainable from the same cells in 2D cultures ^9-12^, suggesting the possibility that cells’ z-dimension could offer one of the distinguishing cues for emerging cell identity. As most 3D culture models involve complex biochemical and biophysical cues, the contribution by cells’ z-dimension to cell identity remains challenging to define.

We explored the molecular players and their modes of action in sensing and regulating cells’ z-dimension by leveraging somatic cell reprogramming into pluripotency, a process that follows choregraphed cell morphological changes. Specifically, reprogramming with the Yamanaka factors begins with somatic cells grown in 2D and end as pluripotent cells in *colonies*. The colony morphology of established pluripotent stem cells (induced pluripotent stem cells, iPSCs; embryonic stem cells, ESCs) is well appreciated and informs the day-to-day assessment of the culture quality: naïve iPSC/ESC colonies are “dome-shaped” and the loss of this domed morphology indicates exit from naïve pluripotency ^13,14^. We report here that the domed colony morphology reflects a minimal cell height of ∼10 μm, resulting from significant nuclear actin accumulation; this morphologic parameter shapes cell identity by tuning ERK activity and subsets of ERK target gene expression.

## Results

### A minimal cell height of 10 **μ**m is required for pluripotency

We utilized somatic cell reprogramming to pluripotency as a model system to investigate the coupling mechanism between cell’s form and identity. Reprogrammable cells with doxycycline (Dox) inducible Oct4/Klf4/Sox2/Myc (OKSM) that co-express GFP reporter from the endogenous *Oct4* locus (R26^rtTA^;Col1a1^4F2A^;Oct4^GFP^) ^15,16^ are used. Shortly after Dox induction (day 4-6), Mouse Embryonic Fibroblasts (MEF) reprogramming cultures contain cells of readily distinguishable morphology: many cells retain fibroblast morphology and small clusters of cells begin to emerge. The fibroblast-appearing cells have a typical height of ∼5 μm (Fig. 1a, Movie S1), on par with the measurements from other cultured mammalian cells ^7,17,18^ (since the nucleus is the largest organelle, nuclear height is often used as an approximation for cell height). In contrast, the cells in clusters reach ∼10+ μm in height (Fig. 1a, Movie S1). As reprogramming progresses, more cells appear in clusters or colonies, some of which begin to activate Oct4:GFP. While the Oct4:GFP-colonies are ∼20 μm in height, the Oct4:GFP+ colonies could reach 40 μm in height (Fig. 1b) consisting multiple cell layers. These results indicate increase in cell height accompanies cell fate transition into pluripotency.

**Fig. 1.**
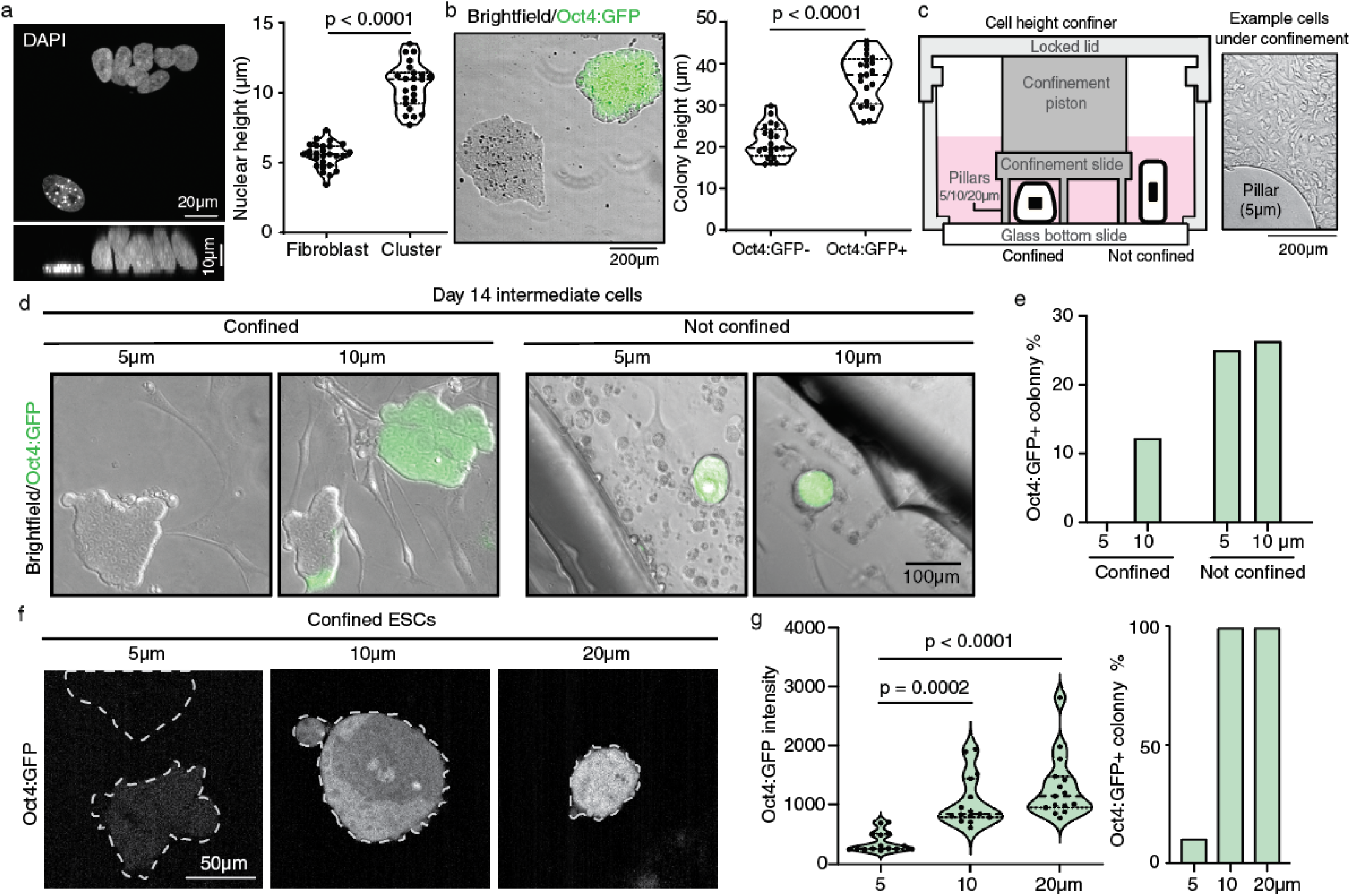
A minimal cell height is required for pluripotency. **a.** Representative top and side view depicting one lone cell (Fibroblast) and a small cluster of cells (Cluster) on reprogramming day 6. Height of DAPI+ nuclei quantified, with median and quartiles shown by thick and thin dashed lines, respectively, on truncated violin plots. n = 27 (Fibroblast), 23 (Cluster). *P* value determined by two-tailed unpaired Student’s t-test. **b.** Representative colony morphology, Oct4:GFP fluorescence and colony height on reprogramming day 14, with median and quartiles shown by thick and thin dashed lines, respectively, on truncated violin plots. n = 21 each. *P* value determined by two-tailed unpaired Student’s t-test. **c.** Schematic of the cell height confiner device, which limits cell height plated underneath by height-defined micropillars. The cells outside of the confinement area share culture medium without height confinement. A representative image of cells grown under 5 μm confinement is shown. **d-e.** Day 14 reprogramming intermediates under 5 or 10 μm confinement. **(d)** Representative brightfield images overlayed with Oct4:GFP fluorescence under 5 or 10 μm confinement, compared with cells outside of the confinement area (not confined). **(e)** Quantification of %Oct4:GFP+ colonies in d. **f-g.** Oct4:GFP+ ESCs under height confinement. Dashed lines delineate colony borders. Oct4:GFP fluorescence of individual optical slices of confocal fluorescence images were used for quantification, with median and quartiles shown by thick and thin dashed lines, respectively, on truncated violin plots, *n* = 15 each. *P* values determined by one-way ANOVA with Kruskal–Wallis tests.

To determine whether a minimal cell height is required for pluripotency, we plated late reprogramming intermediates (day 14) in height confinement devices that limit the maximal height of cells growing underneath by micropillars set to 5, 10 or 20 μm (Fig. 1c). After 16 hours, no Oct4:GFP+ colonies could be found under 5 μm height confinement with the cultures populated by fibroblast-shaped cells and colonies devoid of Oct4:GFP. In contrast, ∼10% colonies under 10 μm confinement were Oct4:GFP+ (Fig. 1d, e). Outside of the confinement area in both devices, similar 20% of the colonies were Oct4:GFP+ which had typical domed morphology (Fig. 1d), indicating that loss of Oct4:GFP occurred specifically to the height-confined cells. These results indicate a minimal cell height is required for pluripotency to emerge. To examine the cell height requirement for pluripotent identity, we plated Oct4:GFP+ ESCs under height confinement. Like the reprogramming intermediates, Oct4:GFP expression in ESCs was contingent on cell height (Fig. 1f, g): under 20 μm confinement, all ESCs remained Oct4:GFP+; 10 μm confinement partially flattened the colonies and reduced Oct4:GFP intensity. Strikingly, all colonies under 5 μm confinement lost Oct4:GFP without any other differentiation-inducing signals (Fig. 1f, g). Taken together, the pluripotent identity can only exist when a specific morphologic requirement is met; inability to reach a minimal height of 10 μm precludes pluripotency. Domed colonies contain tall cells often in multiple layers.

### Actin accumulation inside the nucleus increases nuclear height and promotes pluripotency

Cell morphology is largely determined by the actin cytoskeleton, consisting of dynamically exchanging pools of monomeric G-actin and polymeric F-actin^19,20^. The actin pool is predominantly present in the cytoplasm of most cell types ^21,22^, whereas the existence of nuclear actin has long been considered an impurity or artifact until recently ^23-33^. We observed a significant portion of actin in the nucleus of ESCs and iPSCs (Fig. 2a). To test whether increased nuclear actin allocation accompanies the emergence of pluripotency, we examined the relative abundance of actin in the nuclear and cytoplasmic fractions when large numbers of somatic cells are reprogrammed into Oct4:GFP+ cells, taking advantage of the high reprogramming efficiency of hematopoietic progenitors (Fig. S1a). More actin was found in the nuclear protein fraction as the hematopoietic cells changed into Oct4:GFP+ cells (Fig. 2b, Fig. S1a).

**Fig. 2.**
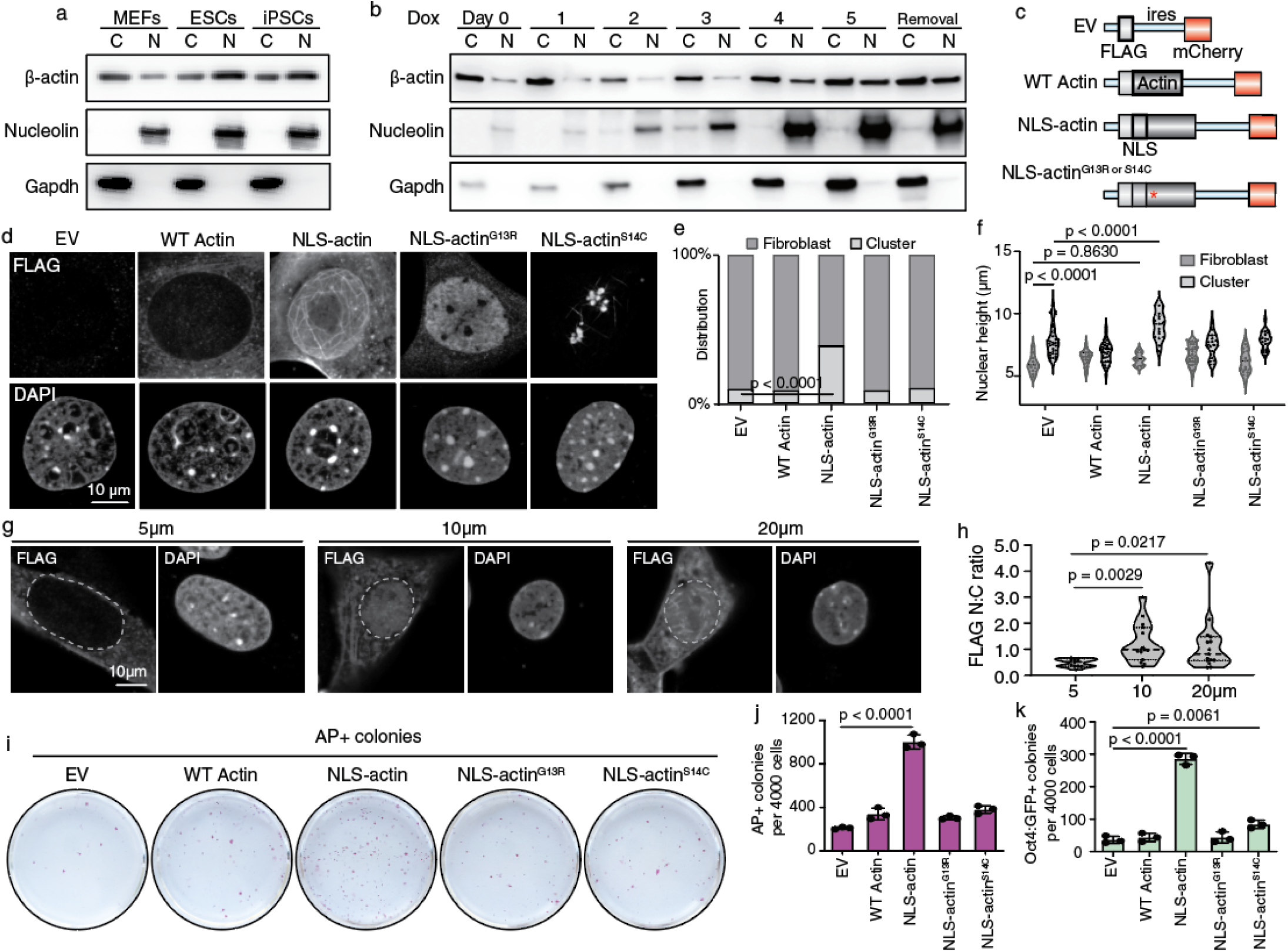
Nuclear actin accumulation increases nuclear height and promotes reprogramming. **a** Western blot for actin in the cytoplasmic (C) and nuclear (N) protein fractions in MEFs, ESCs and iPSCs. Nucleolin and Gapdh control for nuclear and cytoplasmic proteins, respectively. **b** Cytoplasmic and nuclear actin distribution in the hematopoietic cells undergoing reprogramming, sampled at daily intervals for 5 days. Removal: 3 days after Dox removal. **c** Schematic of the retroviral constructs expressing NLS-actin, WT actin and empty vector (EV) control. Red asterisk denotes either of two mutants G13R and S14C. **d** MEFs expressing constructs in c, detected by FLAG immunofluorescence and imaged by confocal microscopy. **e** Frequency of fibroblast-appearing cells and cells in clusters expressing the indicated actin constructs on reprogramming day 6. n = 3 independent replicates per group. *P* values determined by two-way ANOVA with Tukey’s multiple comparisons test. **f** Nuclear height of the cells in each groups of e, with median and quartiles shown by thick and thin dashed lines, respectively, on truncated violin plots. n = 11-35 each (see source data). *P* values determined by two-way ANOVA with Tukey’s multiple comparisons test. **g** FLAG immunofluorescence of NLS-actin+ MEFs under height confinement. **h** Quantification of the nuclear to cytoplasmic (N:C) FLAG signal ratio in cells of g. Nuclear borders are outlined, with median and quartiles shown by thick and thin dashed lines, respectively, on truncated violin plots, n = 11 (5 μm), 15 (10 μm) and 15 (20 μm). *P* values determined by one-way ANOVA with Kruskal–Wallis tests. **i** Representative Alkaline Phosphatase positive (AP+) colonies from reprogrammable MEFs expressing the actin constructs on reprogramming day 10. **j** Quantification of AP+ colonies in i. **k** Quantification of Oct4:GFP+ colonies in i. For j and k, n=3 independent replicates per group. *P* values determined by one-way ANOVA with Dunnett’s multiple comparisons test. Data are presented as mean ± s.d.

To test whether allocating more actin into the nucleus alters cell height, we transduced reprogrammable MEFs with a retroviral construct encoding β-actin, tagged on the N-terminus with a nuclear localization signal (NLS) and a FLAG epitope (Fig. 2c). Empty vector (EV) and wild type (WT) β-actin in identical backbones were used as controls. In cells expressing WT actin, most FLAG signal is in the cytoplasm, confirming actin’s predominant cytoplasmic localization (Fig. 2d). In contrast, FLAG signal is enriched in the nucleus of cells expressing NLS-actin and appeared as elaborate network (Fig. 2d, Movie S2), suggesting that the NLS-actin is involved in F:G dynamics and crosslinked with the endogenous actin. Point mutants NLS-actin^G13R^ and NLS-actin^S14C^, defective in polymerization or depolymerization, respectively ^34^ were also enriched in the nucleus (Fig. 2c, d, Movies S3, S4). As seen before, all reprogramming cultures contained fibroblastic cells and intermediate cell clusters. However, the reprogramming cultures expressing NLS-actin had more cells in clusters (Fig. 2e), and taller nuclei within the clusters (Fig. 2f). The substantial cytoplasmic FLAG signal in the FLAG-NLS-actin+ cells suggested that nuclear actin accumulation could be sensitive to additional cues, i.e. cell height itself. Indeed, NLS-actin fails to accumulate in any cells under 5 μm height confinement (Fig. 2g, h). Thus, the morphologic remodeling typical of early reprogramming reflects increased cell height and nuclear actin accumulation; sufficient cell height is required for nuclear actin accumulation.

We then asked whether nuclear actin accumulation concomitant with cell height increase promotes cell identity change into pluripotency. We found that the NLS-actin expressing cells gave rise to significantly more alkaline phosphatase positive (AP+) and Oct4:GFP+ colonies (Fig.2i-k). In contrast, co-expressing WT actin or NLS-actin^G13R^ made no difference, with NLS-actin^S14C^ having a small increase. The colonies derived from NLS-actin+ cells had sharp borders and domed morphology (Fig. S1b); their resulting Oct4:GFP was brighter on day 10, remained bright, dominated on day 18 and remained Oct4:GFP bright upon Dox removal (Fig. S1c-e), consistent with activation of the *Oct4* distal enhancer ^35^. NLS-actin facilitated reprogramming was also confirmed by wide-spread transcriptomic changes, with day 14 Oct4:GFP+ cells indistinguishable from ESCs (Fig S1f-h). NLS-actin also promoted reprogramming from hematopoietic cells (Fig. S1i-l) whose reprogramming efficiency were already much higher than MEFs (Fig S1a). Lastly, we found that inhibiting the endogenous nuclear actin dynamics by expressing a nuclear localized dominant negative Arp2 (dnArp2) ^36^ inhibited reprogramming (Fig S1m). Taken altogether, these results indicated concentrating F:G competent actin inside the nucleus promotes pluripotency induction in mammalian somatic cells.

### Gtf2i/TFII-I is required for NLS-actin to promote reprogramming

To gain insights into how nuclear actin accumulation and cell height regulate cell identity, we examined the nuclear actin interactome in the intermediate cells by co-immunoprecipitation (Co-IP) followed by mass spectrometry (Fig. 3a-d). We estimated the retrovirally expressed FLAG-NLS-actin to be ∼1% of the endogenous actin, which did not lead to detectible changes in the total actin levels (Fig. 3c, S2a). NLS-actin expression did reduce the protein levels of Sun2 and Lamin A/C (Fig. S2a), lowering of which are molecular features of mechanically soft cells ^37-40^. Silver stain of the nuclear protein precipitates by FLAG antibody from WT actin and NLS-actin expressing cells detected two prominent bands of 42 KD and 250 KD, corresponding to actin and Myh9/10, respectively; these were confirmed by protein-specific antibodies (Fig. 3b-c). Other abundant nuclear proteins such as histone H3 was absent in the precipitated materials (Fig. 3c), confirming the specificity of our approach. A total of 122 proteins with at least 2 unique peptides in the WT actin or NLS-actin samples were recovered (Table S1). These 122 candidates included 38 known actin-binding proteins, 76 known RNA binding proteins and 22 known DNA binding proteins (Fig. 3d). The presence of large numbers of known DNA/RNA binding proteins is consistent with their nuclear-enriched expression.

**Fig. 3.**
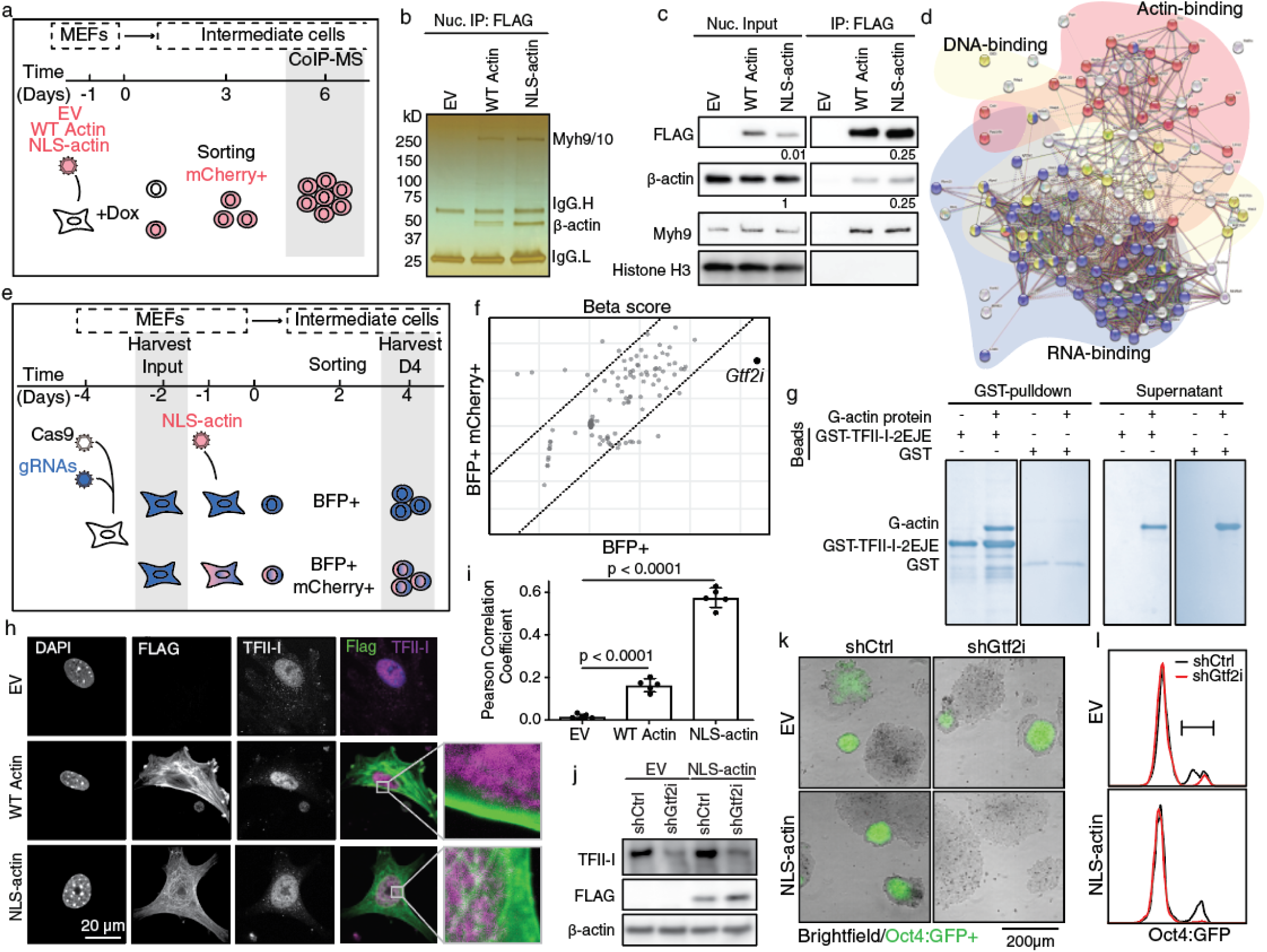
*Gtf2i*/TFII-I is required for NLS-actin to promote reprogramming. **a-d** Co-immunoprecipitation followed by mass spectrometry for identifying nuclear actin interacting proteins. **(a)** Experimental scheme for harvesting proteins for co-IP followed by mass spectrometry. Reprogrammable MEFs expressing EV, WT actin or NLS-actin were reprogrammed for 6 days. Nuclear protein fractions from mCherry+ cells were precipitated by FLAG antibody. **(b)** SDS-PAGE and silver stained nuclear protein immunoprecipitants by FLAG antibody. **(c)** Western blot with antibodies specific for FLAG (exogenous actin) and β-actin (endogenous and exogenous actin), with Myh9 and Histone H3 as positive and negative controls, respectively. Densitometry quantification estimates FLAG+ exogenous actin to be enriched for ∼25-fold; the pulldown accounts for about a quarter of the nuclear input, suggesting overexpression to be ∼1% of that of the endogenous β-actin. **(d)** STRING analysis of 122 candidate nuclear actin interacting proteins containing three categories: actin binding proteins (pink), DNA binding proteins (yellow) and RNA binding proteins (blue). **e-f** CRISPR screen identifying mediator(s) of nuclear actin to promote reprogramming. **(e)** Experimental scheme of the screen. gRNAs library targeting the 122 candidate genes and 10 control genes (4 gRNAs per gene) co-expressing Blue Fluorescence Protein (BFP), and Cas9-expressing vector expressing blasticidin resistance were transduced into reprogrammable MEFs. Input DNA was collected from blasticidin resistant cells before NLS-actin-IRES-mCherry+ transduction. BFP+ cells were sorted into mCherry+/- population and plated separately to continue reprogramming. DNA was collected on reprogramming day 4 from enough cells to provide ∼150x coverage. The choice of this early reprogramming time point was to enrich for regulators of cell morphologic changes and to reduce dominant clone effects in prolonged culture. **(f)** Normalized beta score of BFP+ and BFP+/mCherry+ gRNA reads against input. gRNAs targeting *Gtf2i* are most depleted in BFP+/mCherry+ cells. **g** Recombinant I-repeat domain of TFII-I fused to GST (GST-TFII-I-2EJE), but not GST alone, pulled down purified G-actin. **h** Representative co-IF images of FLAG and TFII-I in MEFs expressing the actin constructs. Inset: zoom in regions across a nuclear boundary region. **i** Quantification of FLAG and TFII-I signal colocalization by Pearson correlation. n=5 independent replicates per group. P values determined by one-way ANOVA with Dunnett’s multiple comparisons test. Data are presented as mean ± s.d. j-l Reprogrammable MEFs expressing either EV or NLS-actin were also transduced with pooled *Gtf2i* shRNAs in reprogramming. **(j)** Confirmation of FLAG-NLS-actin expression and *Gtf2i* knock-down by western blot. (**k)** Representative brightfield and Oct4:GFP colony images on day 15. (**l)** Oct4:GFP FACS on day 12.

To pinpoint the functional mediator of NLS-actin promoted reprogramming, we constructed a guide RNA (gRNA) library targeting the 122 candidate genes. We reasoned that if a specific gene is required for NLS-actin to promote reprogramming, NLS-actin+ cells expressing gRNAs targeting that gene should reprogram less efficiently, resulting in depletion of such gRNAs relative to the others in the library, quantifiable by sequencing. Our library contained 528 gRNAs against the 122 candidates plus 10 control genes, with 4 gRNAs per gene (Table S2). The pooled sgRNA library, co-expressing blue fluorescence protein (BFP), was transduced into Cas9 expressing reprogrammable MEFs (Fig. 3e) and their genomic DNA was harvested as input. BFP+ cells were then transduced with NLS-actin-IRES-mCherry and treated with Dox. Four days after Dox induction, mCherry+ and mCherry-cells were sorted to provide at least 150x coverage, sufficient to detect the reprogramming cells which had an efficiency of ∼7.5% (Fig. 2k). We used FluteRRA (Robust Ranking Analysis) to quantify the abundance of individual gRNAs in the BFP+ and BFP+/mCherry+ cells against control input DNA (Fig. S2b). The gRNAs targeting *Gtf2i* were the most depleted in the BFP+/mCherry+ cells (Fig. 3f), including all 4 individual gRNAs (Fig. 3f, S2c, Table S2). These results indicate that the gene product of *Gtf2i* to be the actin-binding protein responsible for coupling cell morphology and identity.

The *Gtf2i* gene encodes TFII-I, a multifunctional transcription factor containing six I-repeats and is known to regulate signal-induced gene expression ^41-47^. We validated that TFII-I protein is indeed precipitated by NLS-actin in cell lysates (Fig. S2d). AlphaFold ^48,49^ predicts that actin could bind to multiple I-repeat domains on TFII-I ^50,51^ (Fig. S2e, f). We examined these putative interactions by recombinant I-repeat domains fused to GST. Indeed, at least 3 of the 4 I-repeats examined readily pulled down FLAG-NLS-actin from cell lysates (Fig. S2g). GST-I-repeats also pulled down pure actin (Fig. 3g). Furthermore, co-immunofluorescence detected prominent TFII-I signal on the fibrous nuclear actin network (Fig. 3h, i). Taken together, these results demonstrate TFII-I as a novel actin-binding protein.

To independently confirm a functional role of *Gtf2i* in mediating NLS-actin’s pro-reprogramming effect, we designed three shRNAs against *Gtf2i* and confirmed their knockdown in reprogrammable MEFs (Fig. S2h). Of note, *Gtf2i* was previously identified in a genome wide shRNA screen as a mild reprogramming barrier specifically at the transitional stage ^52^. In control cells without NLS-actin expression, *Gtf2i* shRNAs increased colony numbers by ∼2 fold (Fig. S2i-k), similar to the previous report ^52^. By comparison, NLS-actin caused increase in colony numbers was more pronounced. Importantly, all three *Gtf2i* shRNAs decreased NLS-actin-driven increase in colony numbers (Fig. S2h-k). In subsequent experiments, these three shRNAs were pooled together which yielded similar results (Fig. 3j-l). In addition to the increased Oct4:GFP+ colony number, *shGtf2i* led to brighter Oct4:GFP in NLS-actin negative cells (Fig. 3j-l, S2l). In NLS-actin+ cells, however, no Oct4:GFP+ colonies/cells were detected in the presence of *shGtf2i;* the remaining colonies appeared spread or flat (Fig. 3k). Thus, NLS-actin depends on *Gtf2i*/TFII-I to promote reprogramming and to establish the domed colony morphology associated with strong Oct4:GFP expression (Fig. 3k,l).

### The delta isoform of TFII-I, TFII-IΔ, mediates NLS-actin’s pro-reprogramming effect

TFII-I is highly conserved in vertebrates with multiple alternatively spliced isoforms, two of which, β and Δ, are expressed in most cell types including fibroblasts^53,54^. We examined the *Gtf2i* mRNA reads mapping to the alternative region during MEF to iPSC reprogramming and found such reads to be abundant (Fig. 4a, dashed-lined box), suggesting that *Gtf2i* isoform abundance differs across cell states despite similar total *Gtf2i* levels (Fig. S3a). To better resolve the isoforms, we designed qPCR primers spanning the alternative exons (Fig. 4a, black arrows). The transcripts detected by these primers were lower in pluripotent cells, while the relative portion of the *Δ* isoform was much higher (Fig. 4b). Among the reprogramming intermediates (day 6), higher *Δ* was seen in those expressing NLS-actin (Fig. 4c). Consistent with the alternative transcripts level, iPSC expressed a faster-migrating TFII-I protein species, while a larger TFII-I protein band dominated in MEF, corresponding to the Δ and β isoforms, respectively (Fig. 4d). As pluripotent stem cells are mechanically softer ^40,55^, the difference in β and Δ abundance could reflect the isoforms’ differential sensitivity to mechanical cues, although additional means of regulation cannot be excluded. Of note, NLS-actin+ cells had molecular features of mechanical softness (Fig. S2a), suggesting TFII-IΔ’s potential preference to accumulate at low mechanical cues. This notion is further supported by the fact that the Δ isoform dominated when iPSC and ESC were simply plated at high/crowding density (Fig. 4e, S3b). Together, these results implicate TFII-I isoforms to be part of the coupling mechanism between cellular morphomechanics and their identity.

**Fig. 4.**
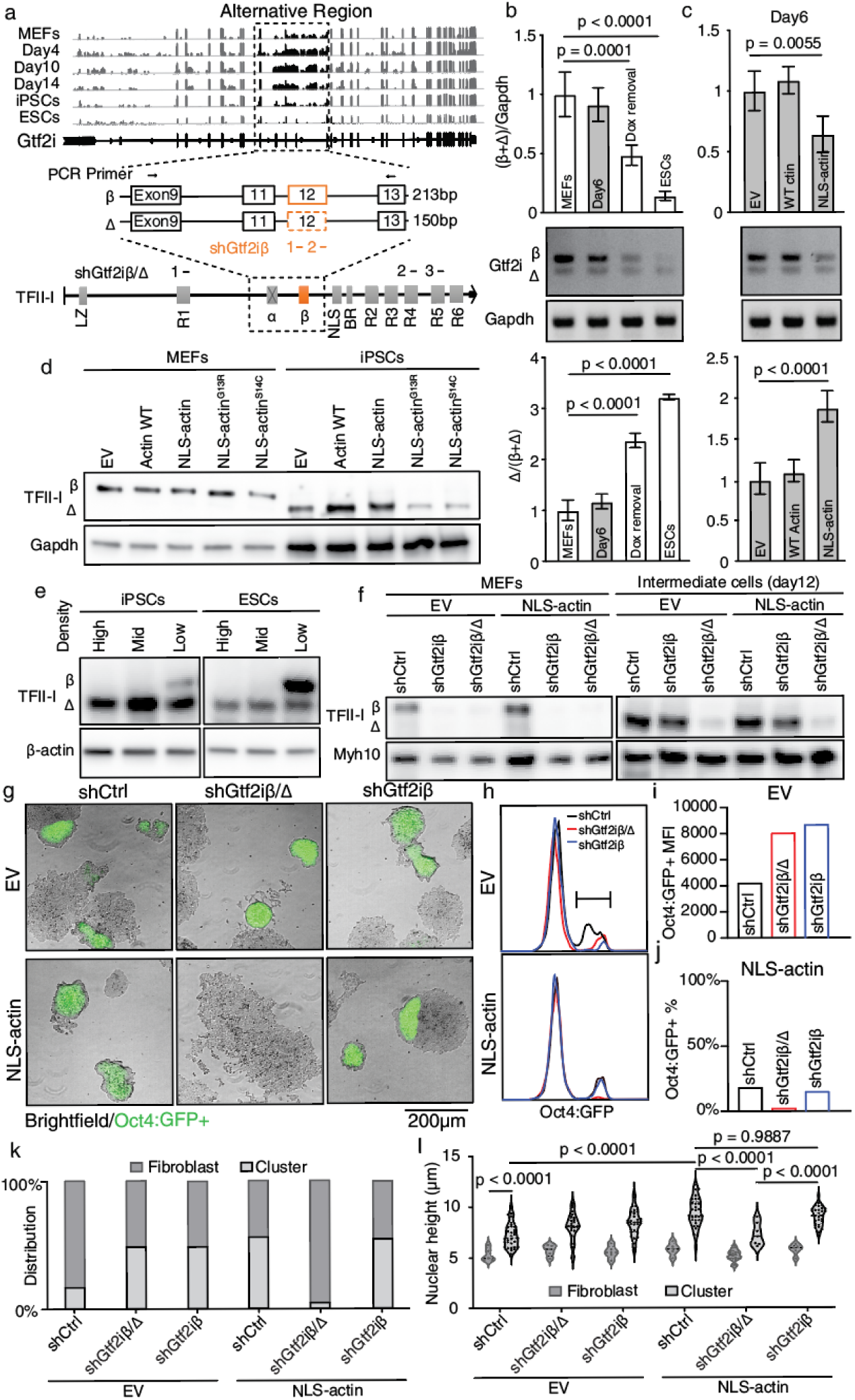
TFII-IΔ mediates NLS-actin’s pro-reprogramming effect. **a** Top: RNA-seq reads mapping to the *Gtf2i* gene during reprogramming. Dashed lined box depicts the region of alternative exons. Middle: Schematic of the alternative exons, highlighting the difference between *β* and *Δ* isoforms. Exon 12 (orange box) is present in *β* but absent in *Δ*. Black arrows above the exons denote positions of qPCR primers, with the anticipated PCR amplicon sizes in base pairs (bp). Bottom: Schematic of TFII-I protein isoforms. The *Δ/β* dual targeting shRNAs are shown as three black bars, with the *β*-specific shRNAs as two orange bars. **b** RT-qPCR in MEFs, day 6 intermediate cells, iPSCs and ESCs, amplifying PCR products corresponding to the *β* and *Δ* isoforms of the *Gtf2i* gene. Quantification of the total product level normalized to *Gapdh* and the *Δ/(β+Δ)* ratio are shown. **c** RT-qPCR detection for *β* and *Δ Gtf2i* in day 6 intermediate cells expressing EV, WT actin or NLS-actin. For (**b**) and (**c**) n=4 each independent replicates per group. *P* values determined by one-way ANOVA with Dunnett’s multiple comparisons test. Data are presented as mean ± s.d. **d** TFII-I western blot in MEFs expressing the actin constructs and the iPSCs derived from the respective MEFs. **e** TFII-I western blot of iPSCs and ESCs plated at different densities. **f** TFII-I western blot of MEFs expressing EV or NLS-actin that also express *β*-specific or *β/Δ* dual targeting shRNAs, and their day 12 reprogramming products. **g** Representative colony morphology and Oct4:GFP fluorescence of cells treated as in f. **h** Oct4:GFP FACS of cells in g. **i** MFI of Oct4:GFP in EV cells. **j** %Oct4:GFP+ in NLS-actin cells. Note the near complete absence of Oct4:GFP+ in NLS-actin cells with *shGtf2iβ/Δ*. **k** Frequency of the fibroblast-appearing cells and cells in clusters in reprogramming cultures. n = 3 independent replicates per group. *P* values determined by two-way ANOVA with Tukey’s multiple comparisons test. **l** Quantification of nuclear height of the cells in k, with median and quartiles shown by thick and thin dashed lines, respectively, on truncated violin plots. n = 5-31 each (see source data). *P* values determined by two-way ANOVA with Tukey’s multiple comparisons test.

To dissect the functional contribution by the TFII-I isoforms, we designed two additional shRNAs targeting *exon 12* (Fig. 4a, two orange bars) present in *β* but absent in *Δ,* while the shRNAs used earlier (Fig. 3j-l, S2h-l) targeted exons common to *Δ* and *β* (Fig. 4a, three black bars) and will be referred to as *Δ/β* dual targeting shRNAs from here on. Either *β*-specific or *Δ/β* dual targeting shRNAs efficiently reduced TFII-I protein in the initial MEFs, which predominantly express the larger β isoform (Fig. 4f, left). In contrast, only the *Δ/β* dual targeting shRNAs reduced the TFII-I protein in later reprogramming cells, which primarily expressed the smaller Δ isoform (Fig. 4f, right; Fig. S3c). Importantly, the *β*-specific shRNAs did not change the number of AP+ or Oct4:GFP+ colonies in NLS-actin expressing cells, contrasting the *Δ/β* dual targeting shRNAs (Fig. S3d-f, compared to Fig. S2i-k). *β*-specific shRNAs also failed to change NLS-actin promoted Oct4:GFP intensity or colony morphology (Fig. 4g-j, compared to Fig. 3k-l, S2l). Consistent with TFII-IΔ in coupling cell form-identity, simultaneous *Δ/β* targeting and NLS-actin expression resulted in flat colonies (Fig. 4g, middle bottom), and flat cells in early reprogramming (Fig 4k,l), an effect absent with *β-*specific KD. These results show that TFII-IΔ participates in the cell form-identity coupling, which we focused on from here on.

### NLS-actin and TFII-IΔ tune ERK activity

The intriguing mode of regulation by NLS-actin and TFII-IΔ suggests additional effector(s) may be at play. TFII-IΔ is an atypical transcription factor, which undergoes cytoplasmic-to-nuclear shuttling upon signaling^37^. Consistent with this notion, much of the TFII-I signal was detected in the cytoplasm following *β-*specific KD (Fig S3c). One of the few known TFII-IΔ effectors is ERK, which can be activated by binding to TFII-IΔ, leading to ERK’s nuclear activation and expression of ERK target genes such as *c-fos* ^41-47^. To determine whether ERK regulation was affected by the NLS-actin-TFII-IΔ relationship, we assessed ERK binding to TFII-IΔ while mutating a critical amnio acid Y248, phosphorylation of which is required for binding ERK ^44^. In reprogramming intermediate cells, the Y248F mutation significantly reduced the binding between phospho-ERK (p-ERK) and TFII-IΔ in the presence of NLS-actin (Fig. 5a). Consistently, while co-expressing WT TFII-IΔ potentiated NLS-actin promoted reprogramming, TFII-IΔ^Y248F^ was ineffective (Fig. 5b; S4a-b). The loss of function by the Y248F mutant was also confirmed by co-expressing mCherry-TFII-IΔ fusion protein, which confirmed the expected changes in Oct4:GFP+ percentage and intensity (Fig 5c, S4c). These results indicate that the ability to bind and regulate ERK to be the mode of action for NLS-actin-TFII-IΔ, corroborating ERK’s well established role in pluripotency^56-63^.

**Fig. 5.**
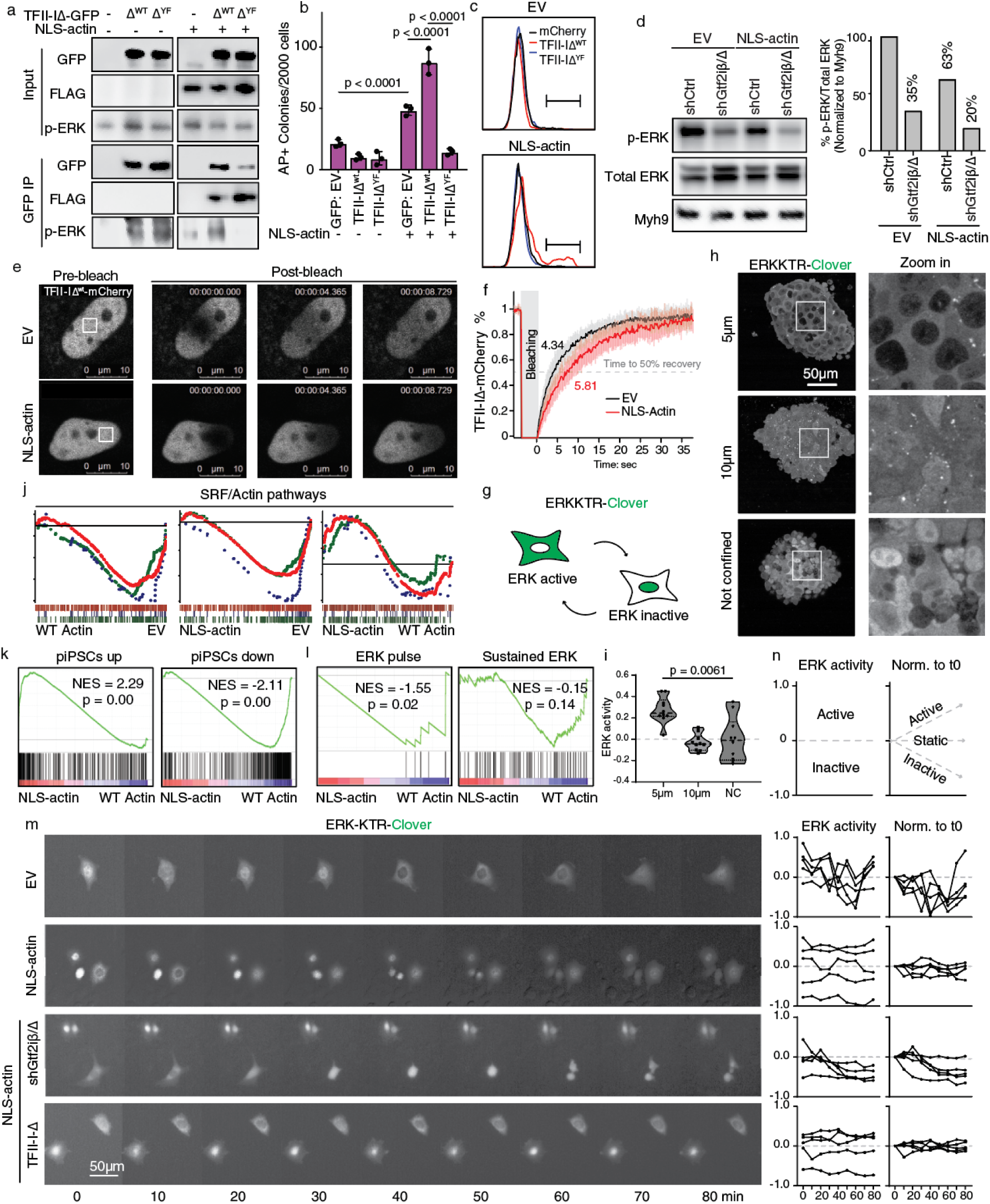
Actin-TFII-IΔ titrate ERK activity. **a** Immunoprecipitation by GFP antibody in cells expressing GFP-tagged TFII-I with or without NLS-actin. **b** Day 10 AP+ colony counts from reprogrammable MEFs expressing the GFP-tagged TFII-IΔ^WT^ or TFII-IΔ^YF^ with or without NLS-actin. n=3 each independent replicates per group. *P* values determined by two-way ANOVA with Tukey’s multiple comparisons test. Data are presented as mean ± s.d. **c** mCherry-tagged TFII-IΔ^WT^ or TFII-IΔ^YF^ performed similarly, measured by Oct4:GFP FACS. **d** Western blot for p-ERK and total ERK in MEFs co-expressing the *β/Δ* dual targeting sh*Gtf2i* with EV or NLS-actin. p-ERK/ERK ratio is quantified and normalized to Myh9. **e** Representative pre- and post-bleach fluorescence of mCherry-tagged TFII-I-Δ. **f** Quantification of the time to half maximal recovery. n=16 each. **g** Schematic of the ERK-KTR reporter. **h** Representative ERK-KTR mClover fluorescence in the reprogramming intermediate cells under height confinement. **i** Quantification of ERK-KTR-mClover fluorescence intensity for cells in h, with median and quartiles shown by thick and thin dashed lines, respectively, on truncated violin plots, n = 10 each independent cells per group, *P* values determined by one-way ANOVA with Kruskal–Wallis tests. **j** GSEA enrichment of three SRF target gene sets in day 4 reprogramming cells co-expressing EV, WT actin and NLS-actin.. **k** GSEA enrichment of gene sets up- or down-regulated in pre-iPSCs. *P* values determined by GSEA. **l** GSEA enrichment of ERK pulse and sustained ERK target gene sets. **m-n** Live cell ERK-KTR-mClover traces in EV, NLS-actin expressing MEFs, with simultaneous *Gtf2i/*TFII-I manipulation. (**m)** Representative time-lapse images of ERK-KTR-mClover. Scale bar =50μm. **(n)** Following the cytoplasmic vs. nuclear KTR intensity over time reveals active, static or inactive ERK. Summary of the traces from five representative cells in m.

We next examined ERK activity by western blotting p-ERK in reprogramming MEFs while manipulating NLS-actin and TFII-IΔ (Fig. 5d). p-ERK was reduced to 35% of the EV control by *shGtf2iΔ/β*, consistent with TFII-IΔ’s role in activating ERK ^47^. p-ERK level was reduced to 63% by NLS-actin expression. Simultaneous expression of NLS-actin and *shGtf2iΔ/β* reduced p-ERK level to 20% of the EV control level. Thus, reprogramming into pluripotency was most efficient when ERK activity was at an intermediate level; further reduction in ERK activity was not conducive. As a reference, we quantified p-ERK levels in MEFs, reprogramming intermediates, iPSCs and ESCs and found much lower p-ERK in pluripotent stem cells, with the intermediate cells transitioning in their p-ERK level (Fig S4d,e). We interpreted these results to mean that *de novo* induction of pluripotency arises within a narrow range of ERK activity; too much or too little ERK activity in the intermediate cells are both counterproductive.

The fact that NLS-actin binds to TFII-IΔ and inhibits the binding between TFII-IΔ and p-ERK (Fig 5a) suggests the possibility that nuclear actin could tune ERK activity by constraining TFII-IΔ’s protein dynamics. To test this, we measured TFII-IΔ protein dynamics by fluorescence recovery after photobleaching (FRAP) ^64,65^ using the TFII-IΔ-mCherry fusion protein. NLS-actin slowed the FRAP kinetics of TFII-IΔ-mCherry (Fig 5e,f), indicating that nuclear actin did reduce TFII-IΔ mobility. As TFII-IΔ activates ERK by cytoplasmic-to-nuclear shuttling, immobilization by nuclear actin would functionally impair ERK activation, explaining why p-ERK was the lowest when *shGtf2iΔ/β* were applied in NLS-actin+ cells (Fig 5d).

To corroborate ERK tuning by actin-TFII-IΔ occurs with cell height change, we imaged ERK activity using a live cell ERK-kinase translocation reporter (ERK-KTR) ^66,67^ in reprogramming intermediates under height confinement (Fig. 5g-i, S4f). The ERK-KTR reporter integrates an ERK docking site (from human Elk1, an ERK substrate), a nuclear import signal (NLS), a nuclear export signal (NES) and mClover. Phosphorylation by activated ERK (p-ERK) favors nuclear export and cytoplasmic accumulation of mClover fluorescence. Conversely, low ERK activity leaves the KTR unphosphorylated, enhancing nuclear mClover fluorescence. Thus, the cytoplasmic to nuclear ratio of mClover fluorescence serves as an indicator of ERK activity in individual cells. At any given time, ERK activity was highest in cells under 5 μm height confinement (Fig. 5h,i), the height that prevented nuclear actin accumulation despite the NLS (Fig 2g,h). Consistent with the previous report, cells under 5 μm height confinement displayed blebbing (Fig S4f) due to increased actomyosin contractility ^7,8^. Importantly, unconfined cells displayed a spectrum of heterogeneous ERK activity (Fig 5i), with cells of 5 μm in height representing the high end of ERK activity spectrum, and cells of 10 μm in height occupying the lower. These results demonstrate that the physiological range of ERK activity can be effected via height modulations around 5-10 μm. Taken together, these results support a model that as cells become taller with actin accumulating in the nucleus, nuclear actin binding to TFII-IΔ constrains the latter’s ability to activate ERK, yielding a cell state with mildly inhibited ERK; cells morphing through different heights explore different ERK activity states.

### Expression of a small subset of ERK target genes respond to tuning by actin-TFII-IΔ

To gain insights into the ERK target genes tuned in this manner, we compared the transcriptomes of reprogramming intermediates expressing NLS-actin, WT actin and EV (Fig. S4g-h). Without Yamanaka factor expression (Vehicle, MEFs), neither actin constructs caused substantial changes as reflected by the small numbers of differentially expressed genes (DEGs) (Fig. S4h, Table S3). 4 days of Dox treatment led to changes in thousands of genes irrespective of the co-expressed actin constructs, as expected from the reprogramming TFs. Co-expression of either WT actin or NLS-actin similarly reduced SRF target genes as assessed by Gene Set Enrichment Analysis (GSEA) ^68,69^, as expected from actin’s known inhibition of MKL1/SRF (Fig. 5j). GSEA also confirmed that day 4 NLS-actin+ cells were more similar to pre-iPSCs (piPSCs as defined by Polo et al. ^70^) than their counterparts expressing WT actin (Fig. 5k). Although NLS-actin+ cells were more advanced toward pluripotency, the DEGs between the actin constructs were rather limited, with 158 up- and 300 down-regulated genes (Fig. S4h). Gene Ontology (GO) analysis could not detect any ERK-related pathways, only identifying “structural constituent of ribosome”, “pre-mRNA intronic binding” and “cadherin-based adhesion” (Table S4).

We therefore hypothesized that the mildly tuned ERK activity could be limited to subsets of ERK target genes, whose activity might be particularly sensitive to short-term ERK activity surges and fluctuations, as cells morphing through different heights (Fig 5h,i; S4f). Subsets of ERK target genes have been defined previously by optogenetically controlled ERK, following either pulsatile or sustained stimulation by light ^71^. We accordingly termed the gene sets responding to these ERK activation regimens as ERK pulse targets and sustained ERK targets, respectively (Fig. 5l). GSEA analysis using these target gene sets show that ERK pulse targets were significantly down-regulated in NLS-actin+ cells, while sustained ERK targets had no significant difference (Fig. 5l). Thus, NLS-actin dampens ERK activity and reduces the expression of a small subset of ERK target genes. Of note, the ERK pulse targets include several immediate early genes known to inhibit pluripotency ^72,73^.

ERK activity is reflected by cycles of activation-inactivation with distinct pulse amplitude and duration according to the signaling cues ^61,71,74-76^. To examine how actin-TFII-IΔ alters ERK pulse behavior, we imaged the live cell ERK-KTR reporter. Tracing the KTR profile over time revealed three types of spontaneous ERK behaviors: activating, inactivating or static (Fig. 5m-n). In control cells (EV), ERK activity was dynamic and heterogeneous, displaying 1-3 activation-inactivation cycles within the observation window (80 minutes). In cells expressing NLS-actin, ERK activity was mostly static. Importantly, however, in cells co-expressing NLS-actin and *shGtf2iβ/Δ*, ERK activity decreased (Fig. 5m-n). Thus, NLS-actin-TFII-IΔ promoted reprogramming is associated with a largely static ERK behavior, with reduced target genes particularly sensitive to ERK pulsatile activities^71^.

### Mild ERK inhibition by chemical inhibitors promotes reprogramming from most fibroblasts

The ERK-tuning model predicts that pluripotency would arise much more frequently if ERK activity is titrated accurately. While cells dynamically exploring their physical environment yields heterogeneous ERK states, a narrow range of ERK activity should be instituted by chemical inhibitors. We therefore treated reprogrammable MEFs with Dox in the presence of 0.05-1 μM PD032591, a well validated MEK inhibitor. Progression toward pluripotency was monitored by %Oct4:GFP+ cells (Fig. 6a). Strikingly, in the presence of 0.1 μM PD032591, >60% of all cells became Oct4:GFP+ by 13 days, and ∼80% cells were Oct4:GFP+ on day 21 (Fig. S5a-b). 2-2.5 fold higher or lower PD032591 at 0.05 and 0.25 μM resulted in substantially lower %Oct4:GFP+. Further increase or decrease in PD032591 concentrations all reduced %Oct4:GFP+ to the level of DMSO control. Because 1 μM PD032591 is the concentration in the 2i media for cultivating naïve pluripotent stem cells ^60^, we assessed the extent of ERK inhibition at 0.1 μM PD032591 by p-ERK western (Fig. 6b,c). Of note, p-ERK levels dropped precipitously between reprogramming day 4 and 7. When p-ERK level was high on day 4, the inhibition by 0.1 μM PD0325901 was barely detectible; however, inhibition of p-ERK by 0.1 μM PD0325901 became clear on day 7 (29% of DMSO). Similar to PD0325901, another ERK inhibitor, U0126 also significantly increased %Oct4:GFP+ cells at low dose, albeit to a lesser extent than PD0325901 (Fig. S5c-f). The range of ERK inhibition resulting from the low dose inhibitor treatments is in good agreement with that achieved by NLS-actin expression or *shGtf2iβ/Δ,* where reducing p-ERK to below 20% of the control level became counterproductive to reprogramming as seen in cells simultaneously expressing NLS-actin and *shGtf2iβ/Δ* (Fig 5d).

**Fig. 6.**
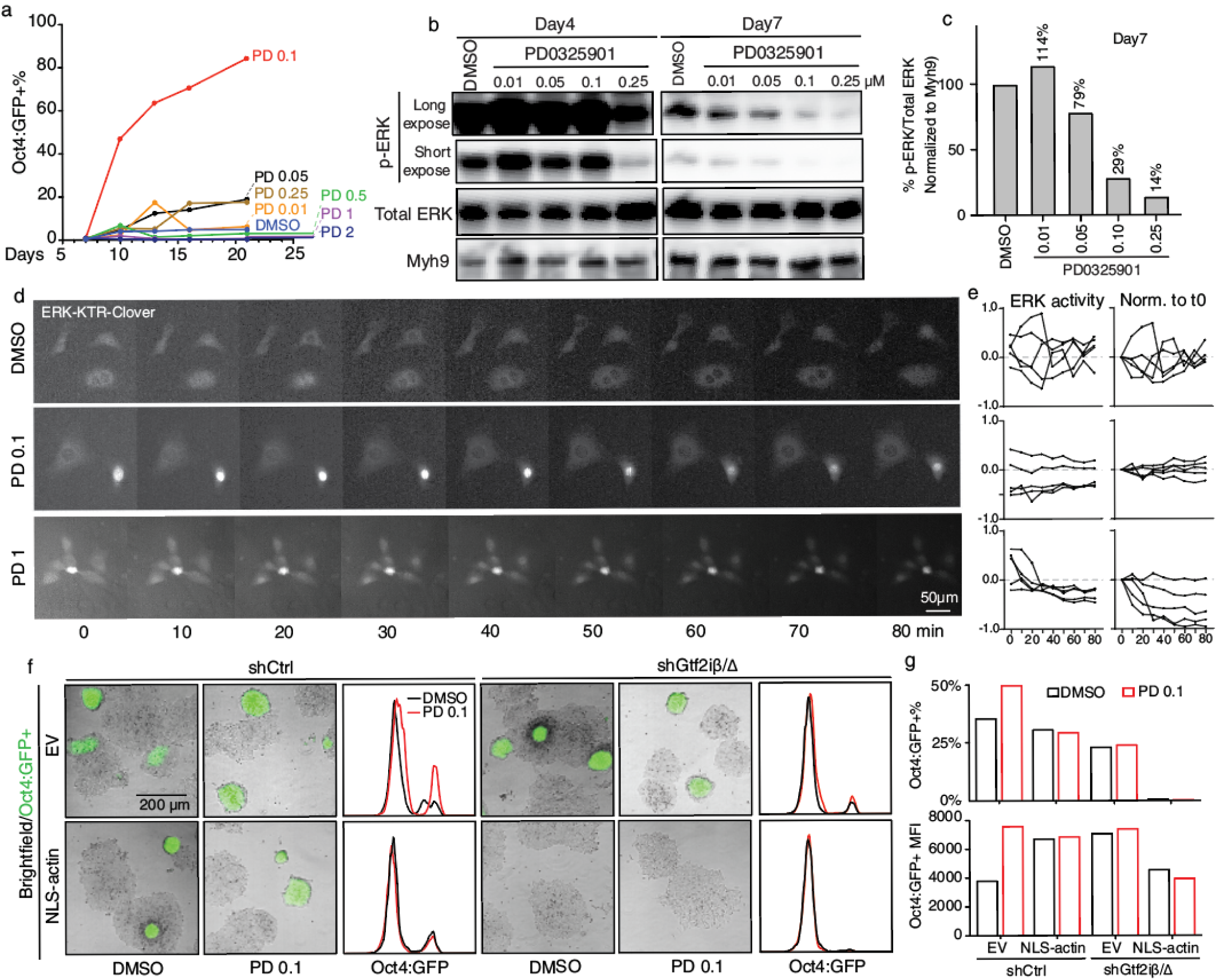
Mild ERK inhibition by chemical inhibitors recapitulates morphomechanic tuning and promotes reprogramming. **a** %Oct4:GFP+ cells arising in reprogramming cultures treated with 0.01 μM to 2 μM of PD0325901 over time (days), as determined by FACS. **b-c** MEFs on reprogramming day 4 and 7 in the presence of various PD0325901 concentrations. **(b)** Representative p-ERK/ERK western blots with Myh9 as loading control. (**c)** Quantification of p-ERK/ERK ratio on day 7, normalized to Myh9. **d-e** Single cell ERK-KTR-mClover traces of MEFs in low (0.1 μM) and high (1 μM) concentrations of PD0325901. **(d)** Representative time-lapse images of ERK-KTR-mClover. Scale bar = 50μm. **(e)** Quantification of cells similar in d. Summary of the traces from five representative cells in d. **f** Representative bright field/Oct4:GFP fluorescence in cells with actin-TFII-I manipulation, in the presence of 0.1 μM PD0325901 (PD 0.1). **g** Quantification of %Oct4:GFP and MFI for cells in f.

We next examined the live ERK activity in cells treated with low dose inhibitors (Fig. 6d-e). In DMSO treated control cells, ERK was dynamic, displaying 1-3 active-inactive cycles as before (Fig. 6d-e, top). In the presence of 0.1 μM PD0325901, ERK became largely static (Fig. 6d-e, middle), similar to cells expressing NLS-actin or *shGtf2iβ/Δ* (Fig. 5m-n). In 1 μM PD0325901, ERK activity decreased (Fig. 6d-e, bottom), resembling the cells simultaneously expressing NLS-actin and *shGtf2iβ/Δ* (Fig. 5m-n, third row). The effect of low dose PD0325901 was also recapitulated in the brightness of Oct4:GFP (Fig 6f,g): while 0.1 μM PD0325901 increased the Oct4:GFP intensity in control/EV cultures (Fig 6f, top left), it was ineffective in cells that already express NLS-actin or *shGtf2iβ/Δ* or both, further supporting that the emerging Oct4:GFP+ identity is contingent on a very specific ERK activity level. Taken together, while somatic cell reprogramming into pluripotency usually occurs infrequently, this rare change in cell identity is greatly facilitated when ERK activity is titrated within a narrow range (Fig. S5g).

## Discussion

We propose that cell morphology regulates identity by interpreting ERK activity in a finely tuned manner; this is accomplished by pooling actin in the nucleus where actin binding to TFII-IΔ constrains ERK activation. Our earlier work revealed that reprogramming stochasticity is not experienced by rare cells of ultrafast cell cycle^77,78^, and their prospective isolation enriched cells of a much less spread morphology ^3,79,80^, which we show in this report to be taller. As cell morphology is influenced by physical parameters ill-controlled in most routine cultures, the apparent stochasticity of Yamanaka reprogramming could be appreciated as the chance encounter with a narrow ERK activity level permissive for this fate change (Fig. S5g) ^81-84^. As the components of the actin-TFII-IΔ-ERK axis are ubiquitous to all cells, this morphomechanic tuning mechanism likely operates beyond reprogramming, e.g. when cells exit pluripotency or undergo epithelial to mesenchymal transition (EMT), both of which experience pervasive morphological remodeling. Spontaneously differentiating cells in naïve pluripotency cultures are identified by flattening colonies, likely representing cells of increased ERK activity. In this regard, ERK activation has also been reported to occur through decreased membrane tension^63,85^. ERK activity could gain by higher pulsing amplitude, frequency and/or duration ^61,71,74-76^. While the tuning by actin-TFII-IΔ led to apparently fewer and/or lower amplitude in the spontaneous ERK pulses (Fig 5m,n), tracking ERK pulses as cells undergo cell fate transitions guided by developmental signals would yield deeper understanding for how morphomechanics consolidate identity, especially in 3D culture models where authentic cell identities emerge ^9-12^. Lastly, as genetic inactivation of ERK or prolonged cultivation in 1 μM PD032591 impair stem cell quality ^58,59,86-88^, it remains to be determined whether and how finely tuned ERK improves stem cell quality.

Nuclear actin is known to be abundant in frog oocyte germinal vesicles ^89^. These exceptionally large cells collapse due to gravity when the nuclear actin meshwork is disrupted ^90^. Therefore, it is perhaps not surprising why polymerized actin is required in the germinal vesicle to reprogram transplanted somatic nuclei, shown by Gurdon and colleagues ^91^. Mammalian somatic cells are orders of magnitude smaller, where the possibility that nuclear actin primarily functions by mechanical support becomes tenuous. However, the much smaller somatic nuclei do undergo prominent “swelling” in this system ^92^. Filamentous actin also form in mammalian somatic nuclei at mitotic exit to expand the compact postmitotic nuclei to that of G1 conformation^93^. Across these diverse biological contexts, a coherent theme appears to be nuclear actin’s role in modulating the nuclear size. In this regard, we found that nuclear actin content is directly related to the nuclei’s z dimension.

How nuclear dimension regulates cell behavior has only begun to be appreciated. Addressing such questions requires approaches for accurately manipulating the nuclear dimensions while assessing cellular behavior, which has so far been largely conducted on short time scales ^7,94,8^. In the 3D cultures that do faithfully recapitulate the varieties of cell identity, it remains challenging to tease apart the contribution from nuclear dimension changes versus other cellular, chemical and physical cues^95-97^. Our work builds on the otherwise identical culture condition for reprogramming, only applying height restrictions to the reprogramming cells. For cells grown in 2D, confining their height to 5 μm, but not 10 μm, stretches and activates nuclear membrane/endoplasmic reticulum-localized mechanosensitive channels leading to Ca2+ directed actomyosin contractility, as evidenced by extensive blebbing ^78^. Insights from the short time scale studies paint a model for how limiting nuclear dimension enables cells to escape the immediate confinement; their activated contractility could also explain why actin fails to concentrate in flattened cells despite the extra NLS (Fig. 2g-h), as contractility likely draws actin away from the nucleus ^7^. Consistent with flat cells being more contractile, blebbing indeed occur in reprogramming cells of 5 μm height (Fig S4f). In this regard, taller cells could curtail contractility by stowing actin away from the cytoplasm while buffering mechanical stretching by having their nuclear membrane less taut. The full picture for morphomechanic cell fate control is likely much more complex, as it plays out on the time scale of development and tissue homeostasis, and as ERK-actomyosin contractility engage in complex relationships ^98 99,100^. However, as recent studies reveal critical roles for nuclear actin in DNA repair^101,102^ and RNA polymerase bursting ^103^, it would be informative to begin assessing how nuclear dimension regulates these fundamental processes by controlling the nuclear actin pool sizes, potentially also contributing to pathology^104^.

Our approach in identifying actin-binding proteins is inherently biased toward abundant nuclear proteins and/or those with high binding affinity, nominating TFII-I’s novel biology. In mice, *Gtf2i* inactivation results in early embryonic lethality ^105^. In humans, hemizygous deletion of *GTF2I* genomic region is associated with neurodevelopmental deficits known as Williams-Beuren Syndrome, while its duplication leads to autism spectrum disorders ^105-107^. Single nucleotide polymorphisms at *GTF2I* loci are associated with autoimmune diseases ^108-110^. A point mutation (L424H) is prevalent in thymic epithelial tumors ^111^. Contrasting its importance in development and diseases, the understanding of how *GTF2I* abnormality causes diseases is limited. Our work suggests that pathogenesis by TFII-I could occur via altered ERK tuning^46,112,42^, as the interfaces along the actin-TFII-IΔ-ERK axis could be modified by mutations. How TFII-I-ERK precisely regulates the expression of cell identity gene awaits further investigation.

## Material and Methods

### Cell culture and reprogramming

All mouse work was approved by the Institutional Animal Care and Use Committee of Yale University. The reprogrammable mice with reporter (R26rtTA;Col1a14F2A;Oct4GFP) were derived by crossing reprogrammable mice with Oct4:GFP mice, which has been described before.

### DNA constructs

All actin constructs were cloned into pMSCV-IRES-blasticidin or pMSCV-IRES-mCherry backbone. The shRNAs targeting *Gtf2i* and its *β* isoform were generated by inserting the short hairpin sequence into the lentiviral backbone psi-LVRU6MP (GeneCopia). All sequences are listed in Supplementary Table 5. The pSFG-GFP, pSFG-TFII-I-GFP delta and pSFG-TFII-I-GFP Y248&249F were from Addgene (#22199, #22190, #22196), and the TFII-I-mCherry fusion proteins are also cloned into pMSCV-IRES-mCherry backbone. The plasmids GST-TFII-I-2D9B, GST-TFII-I-2DN4, GST-TFII-I-2ED2 and GST-TFII-I-2EJE were generated using the pGEX6P vector as the backbone. The ERK-KTR-Clover is from Addgene (#90227).

### Cell Height Confinement

6-well static cell confiner device (4Dcell, France) was used to confine cells at different heights according to the manufacturer’s instructions. The confinement cover slides used in this study contains 5, 10, 20µm pillars. For Oct4:GFP colony counting and Oct4:GFP intensity measurement, reprogramming intermediate cells or ESCs were seeded in 6-well plates, and height confinement was applied overnight. ImageExpress Micro 4 Imaging system or Leica Stellaris confocal microscope platform were used to count Oct4:GFP colony and measure Oct4:GFP intensity, with live cell environmental control at 37°C, compressed gas of 5% CO_2_, 21% O_2_ and balance Nitrogen. For FLAG staining, NLS-actin transduced MEFs and day 12 reprogramming cells were fixed after height confinement. N/C ratio of FLAG intensity was calculated using LAS AF software. For imaging ERK-KTR Clover under height confinement, cells were confined for 3 hours.

### Western blotting and immunofluorescence

All procedures and antibodies used in protein analyses are listed in the accompanying supplementary materials.

### Construction of custom sgRNA library and screening

The online web tool CHOPCHOP (https://chopchop.cbu.uib.no/) was used to generate sgRNA designs against target genes. For each gene, 4 sgRNAs were chosen based on the location and score. Screening is done by following the Zhang Lab’s protocols with minor modification.

### ColabFold Modeling and GST-pulldown

ColabFold^49^, a access tool of protein structure and complex prediction using MMseqs2 of AlphaFold^48^ was used to modeling interaction of *Actin* and *Gtf2i* or its known structured repeat domains *1q60*. All protein/domain sequences were obtained from Uniprot. All default parameters “msa_mode”: “MMseqs2 (UniRef+Environmental)”, “model_type”: “AlphaFold2-multimer”. To model the *Actin* and *Gtf2i*, residue boundaries for the *Gtf2i* were 1-979AA, and *Actin* were 980-1354AA. To model the *Actin* and *1q60* complex, residue boundaries for the *1q60* were 1-99AA, and *Actin* were 100-474AA.

### RNAseq and analysis (GSEA, CellNet)

RNA-seq libraries were prepared with TruSeq Stranded mRNA Library Prep Kit (Illumina, RS-122-2101) following the manufacturer’s instructions. Sequencing was performed with the Illumina HiSeq 4000 Sequencing System. For data analysis, the RNA-seq reads were mapped to mouse genome (mm10) with TopHat2 software. Gene abundance was calculated using cuffnorm, with gene expression levels and Fragments per kilobase per million (FPKM). Genes with FPKM ≥1 in two or more samples were selected for further analysis. Differentially expressed genes (DEGs) were identified by Cuffdiff followed by cutting off with FDR-adjusted P value <0.05 and fold change >2. MA plot of differentially expressed genes was also done with the R software. RNA-seq raw data and processed data have been deposited as GSE229191. GO analysis of differentially expressed genes was performed with R.

### Supplemental Materials (with full description of materials and procedures)

Four supplementary tables (Table S1-4) and movies (Movie S1-4) accompany this manuscript. Supplementary Table S5 contains sequences for all primers used.

## Acknowledgements

We thank the members of the Shangqin Guo laboratory and Jun Lu laboratory for critical input and discussions.

## Contributions

Q.W designed, performed, analyzed most of the experiments, produced figures and tables, and wrote the manuscript. J.Z. performed bioinformatics analysis. B.L performed statistical analysis and produced figures. X.H, B.M.F, S.M.S, K.K provide mouse maintenance, MEF cell generation and intellectual support.

W.C and E.M.D.L.C. provided purified actin proteins and GST pull down support.

## Corresponding author

Correspondence to Shangqin Guo.

## Ethics declarations

Competing interests

All authors declare no competing interests.

---

**Fig. S1.**
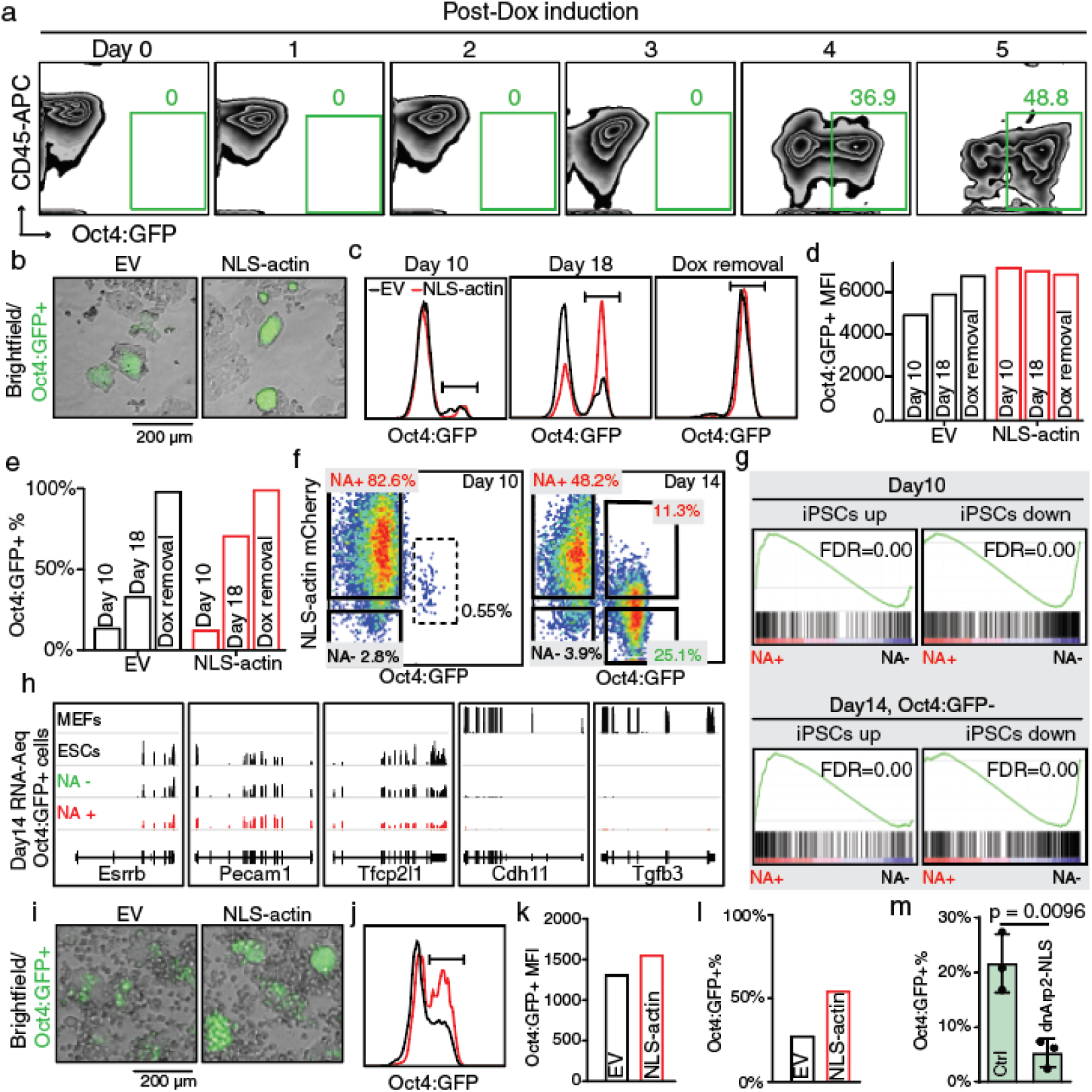
NLS-actin promotes reprogramming. **a** Reprogrammable hematopoietic progenitors undergoing reprogramming and analyzed by FACS for CD45 (hematopoietic marker) and Oct4:GFP at daily intervals for 5 days. **b** Representative colony morphology from reprogrammable MEFs expressing EV or NLS-actin. **c** Oct4:GFP FACS for cells in b. **d** The Mean Fluorescence Intensity (MFI) of Oct4:GFP in c. **e** %Oct4:GFP+ cells in b. **f** Representative FACS plots for reprogramming MEF cultures expressing NLS-actin-IRES-mCherry on day10 and day14. The gated populations were FACS sorted for RNA-Seq analysis. **g** Gene Set Enrichment Analysis (GSEA) with previously defined gene sets (Polo et al.) between cell populations sorted from f. **h** Expression of representative naive pluripotency genes (*Esrrb, Pecam1 Tfcp2l1*) and fibroblast genes (*Cdh11, Tgfb3*) in the Oct4:GFP+ cells sorted from f (day 14). **i** Representative colony morphology from reprogrammable hematopoietic progenitors expressing EV or NLS-actin. **j-l** Representative FACS analysis of Oct4:GFP for cells in i. **m** MFI and % of Oct4:GFP for cells in i**. m** %Oct4:GFP+ cells from reprogrammable MEFs expressing control vector (Ctrl) or dominant negative (dn)Arp2-NLS on day 12. n=3 independent replicates per group. *P* values determined by unpaired *t*-test with Welch’s correction. Data are presented as mean ± s.d.

**Fig. S2.**
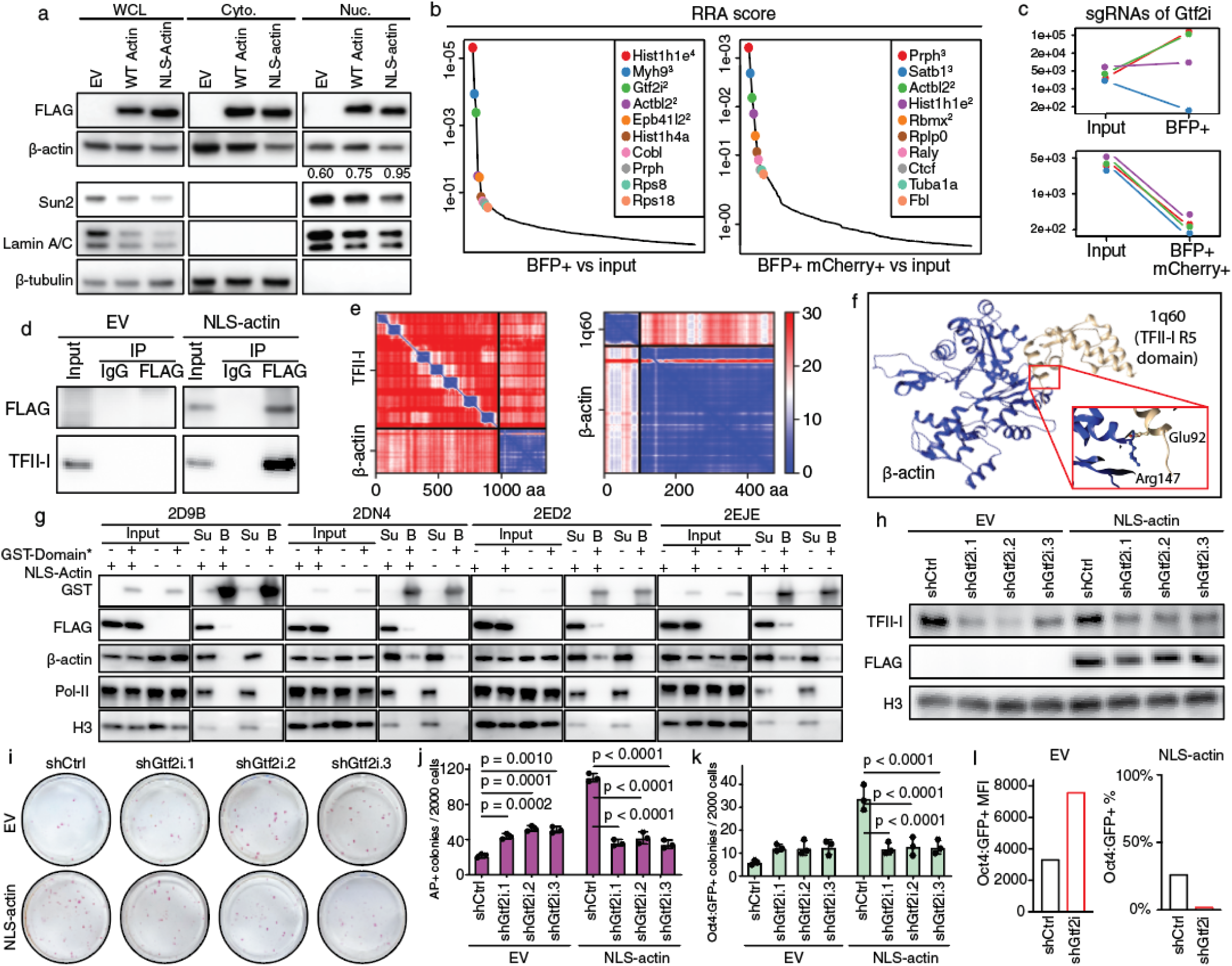
*Gtf2i*/TFII-I is required for NLS-actin to promote reprogramming. **a** Similar to Fig 3a-c. Western blot of whole cell lysates (WCL), cytoplasmic and nuclear lysates from reprogrammable MEFs expressing EV, WT actin or NLS-actin on day 6. Sun2 and Lamin A/C are two nuclear protein controls and β-tubulin is cytoplasmic control. The numbers denote the ratio between nuclear and cytoplasmic actin band intensity. **b-c** CRISPR screen analysis using MAGeCK and MAGeCKFlute tools. **(b)** FluteRRA (Robust Ranking Analysis) of gRNAs in BFP+ or BFP+/mCherry+ cells. Superscript denotes the number of enriched gRNAs. **(c)** Absolute reads number of the four individual gRNAs targeting *Gtf2i* in BFP+ and BFP+/mCherry+ cells. **d** Western blot for endogenous TFII-I in the FLAG IP products. **e** AlphaFold2 prediction of the interaction between full length TFII-I (aa1-979) and β-actin (aa980-1354), or between one of the I-repeat domains (1q60, aa1-99) and β-actin (aa100-474). **f** AlphaFold2 prediction of the interaction between actin (blue) and one of the I repeat domains of TFII-I1q60 (yellow) implicating Arg147 in actin and Glu92 in 1q60. **g** GST-pull down of FLAG-tagged actin by GST tagged TFII-I I-repeat domains 2D9B, 2DN4, 2ED2 and 2EJE. Su: protein in unbounded supernatant; B: precipitates by GST-beads. Pol-II and H3 included as negative controls. **h-k** Reprogrammable MEFs expressing EV or NLS-actin, transduced with three individuals *Gtf2i* shRNA in reprogramming. **(h)** Confirmation of FLAG-NLS-actin expression and *Gtf2i* knock-down by western blot, with Histone H3 as a loading control. **(i)** Representative AP+ colonies on day 10 from cells in h. **(j)** Quantification of AP+ colonies in i, **(k)** Quantification of Oct4:GFP+ colonies in i. For (**j**) and (**k**) n=3 each independent replicates per group. *P* values determined by two-way ANOVA with Dunnett’s multiple comparisons test. Data are presented as mean ± s.d. **l** Quantification of Oct4:GFP MFI and %Oct4:GFP+ for cells in Fig 3l. MFI was not quantified in NLS-actin+ cells due to their absence.

**Fig. S3.**
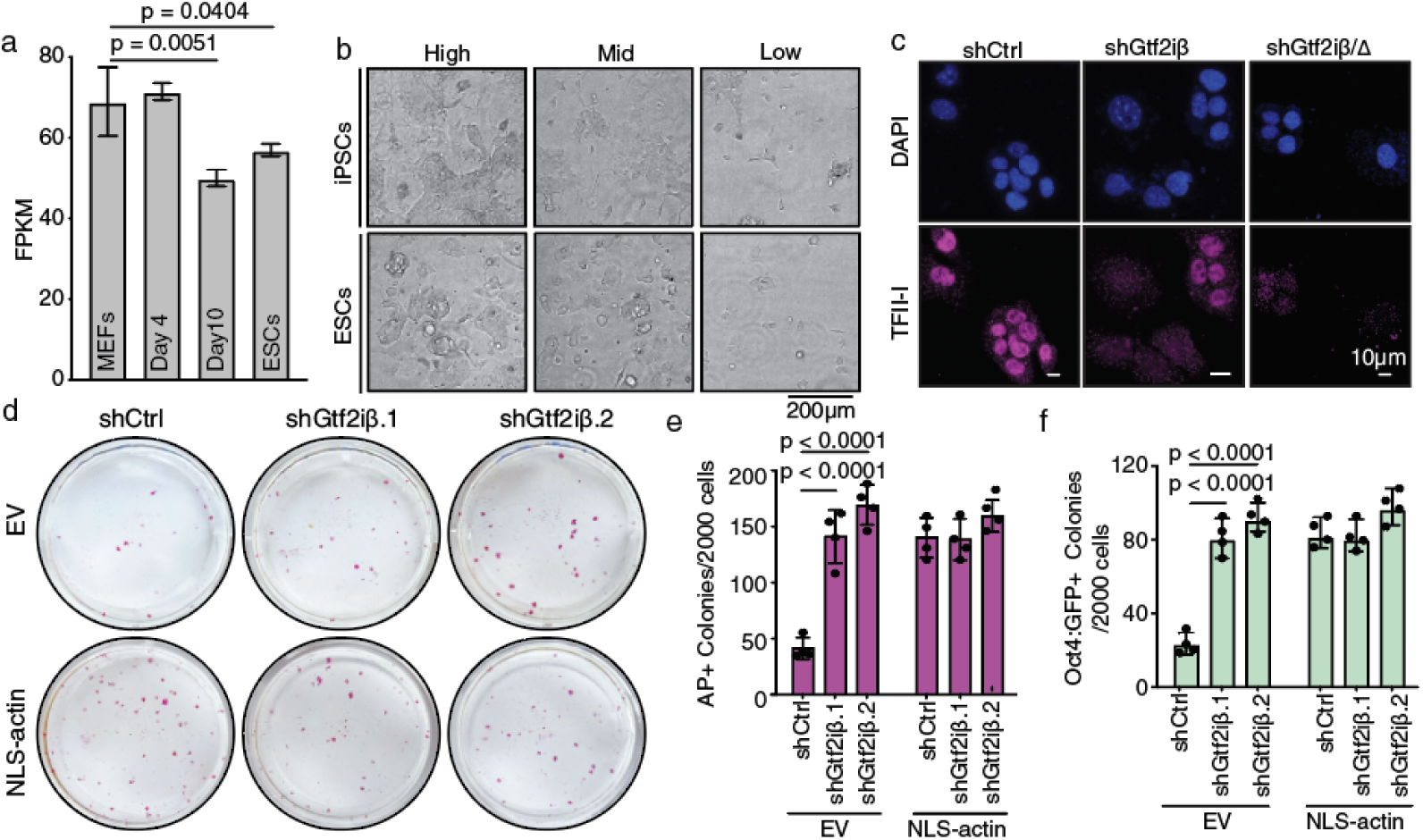
TFII-IΔ mediates NLS-actin’s pro-reprogramming effect. **a** Total FPKM mapping to *Gtf2i* in various cell types. n=2 for MEFs and Day4, n = 3 for Day10 and ESCs, *P* values determined by two-way ANOVA with Dunnett’s multiple comparisons test. Data are presented as mean ± s.d. **b** Representative iPSCs and ESCs plated at different densities for western blotting in Fig 4e. **c** TFII-I immunofluorescence in reprogramming intermediate cells in the presence of *Gtf2i* shRNAs. **d** Representative AP+ colony images (day 10) from reprogrammable MEFs expressing either EV or NLS-actin, combined with *Gtf2iβ-*specific shRNAs. **e** Quantification of AP+ colonies in d. **f** Quantification of Oct4:GFP+ colonies from cells similar to d. For (**e**) and (**f**) n=4 each independent replicates per group. *P* values determined by two-way ANOVA with Dunnett’s multiple comparisons test. Data are presented as mean ± s.d.

**Fig. S4.**
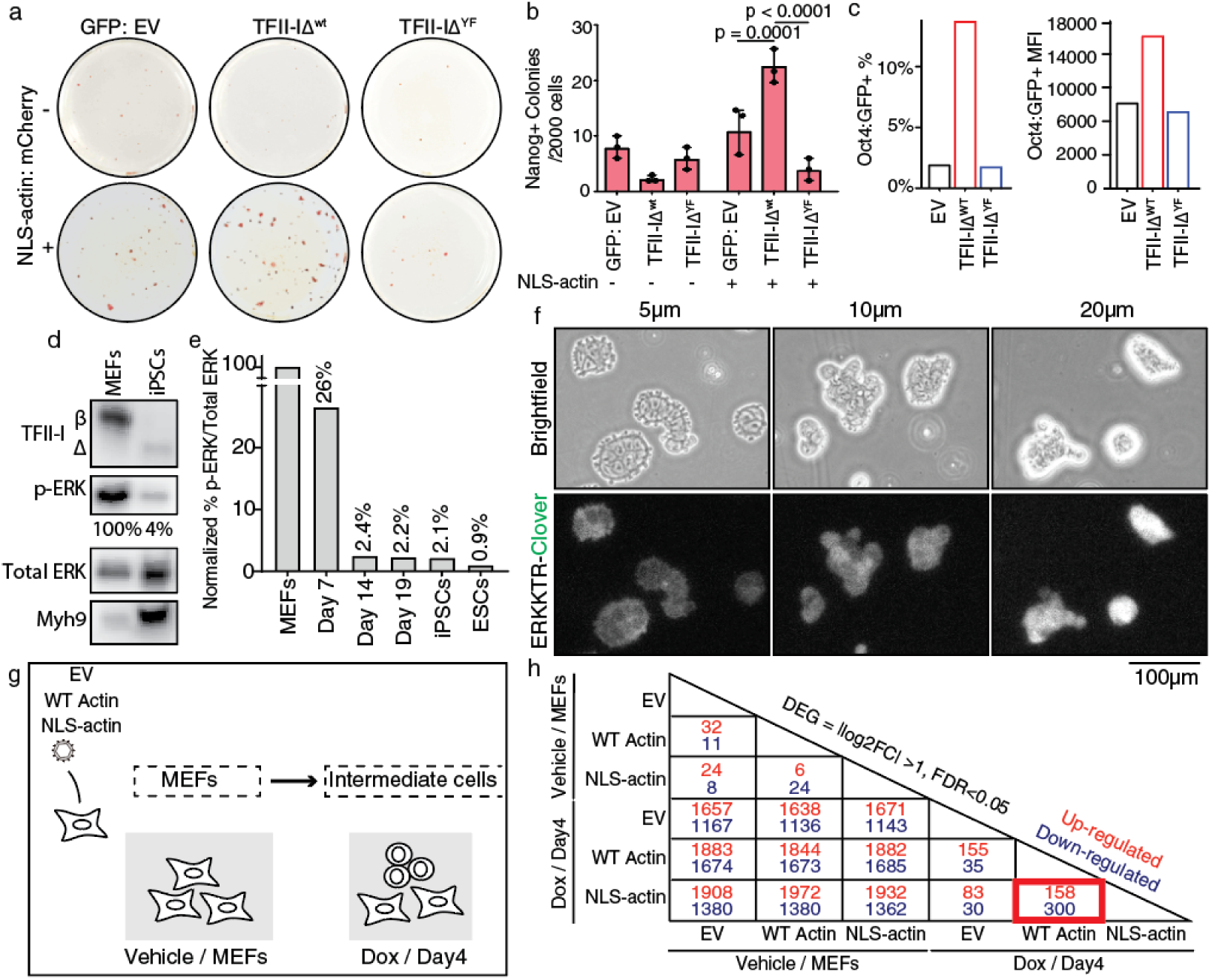
Actin-TFII-IΔ titrate ERK activity. **a** Representative day 10 AP+ colonies from reprogrammable MEFs expressing the GFP-tagged TFII-IΔ^WT^ or TFII-IΔ^YF^ with or without NLS-actin co-expression. **b** Quantification of Nanog+ colonies with cells shown in a, n=3 each independent replicates per group. *P* values determined by two-way ANOVA with Tukey’s multiple comparisons test. Data are presented as mean ± s.d. Related to Fig 5b. **c** %Oct4:GFP+ and Oct4:GFP MFI quantification in cells expressing the mCherry-tagged TFII-IΔ^WT^ or the Y248F (TFII-IΔ^YF^) mutant, related to Fig. 5c. **d** Western blot for TFII-I and p-ERK/ERK in MEF and iPSCs. **e** Quantification of p-ERK/total ERK in MEFs, intermediate cells, iPSCs and ESCs following similar approaches shown in d. **f** Representative brightfield and ERK-KTR-mClover images of the reprogramming intermediate cells under height confinement. Note the prominent blebbing from cells under 5 μm confinement. **g** Experimental workflow for harvesting cells for RNA-seq analysis. Reprogrammable MEFs transduced with EV, WT actin or NLS-actin were cultured in either vehicle or Dox for 4 days. **h** DEGs in pair-wise comparisons across all sample groups.

**Fig. S5.**
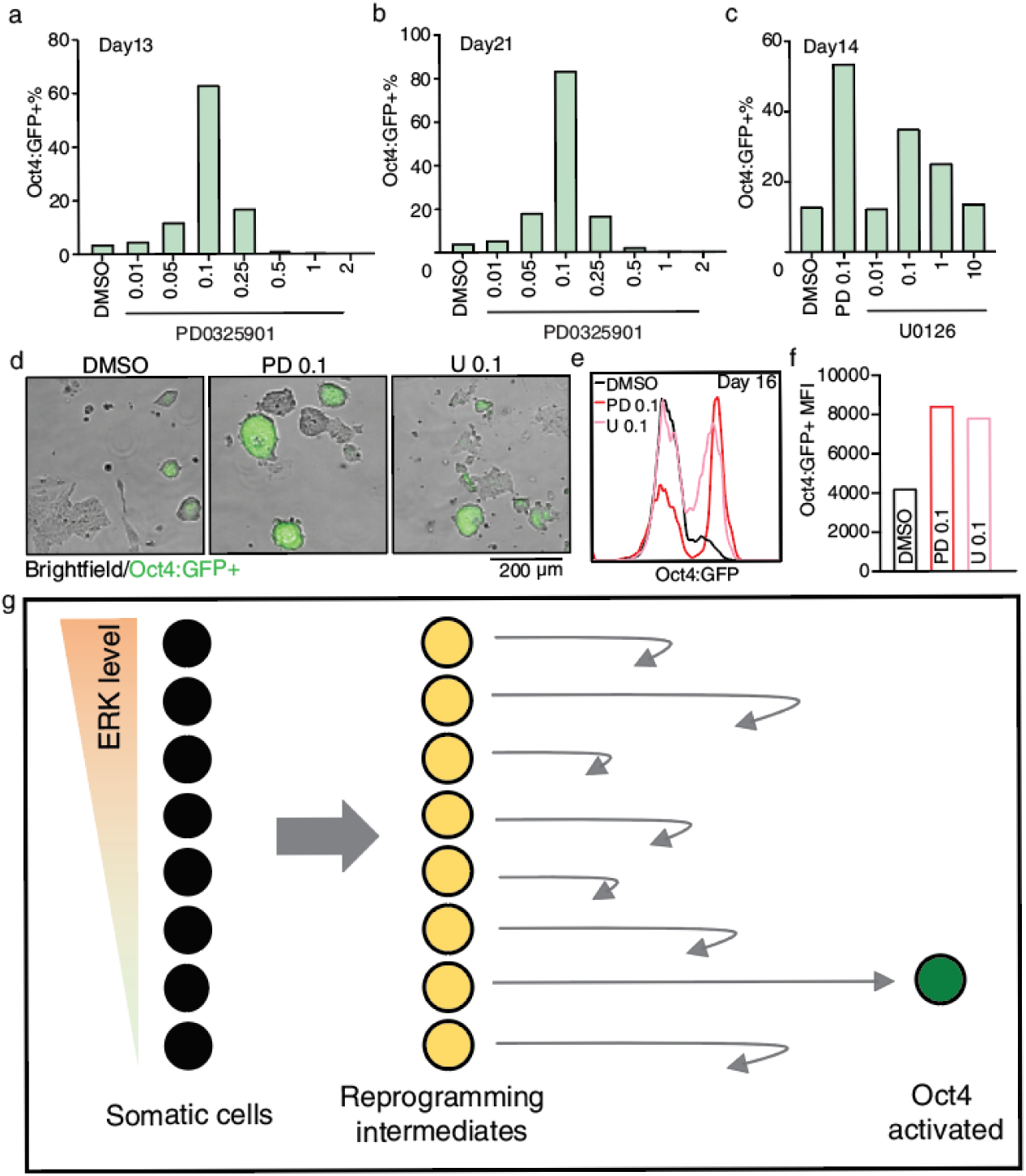
Mild ERK inhibition by chemical inhibitors promotes reprogramming. **a-b** %Oct4:GFP+ cells in reprogramming cultures treated with different concentrations of PD0325901 on **(a)** day 13 and **(b)** day 21. Related to Fig. 6a. **c-f** Reprogramming cultures treated with different concentrations of U0126. **(c)** %Oct4:GFP+ cells on day 16. 0.1 μM PD0325901 included as positive control. **(d)** Representative bright field and Oct4:GFP fluorescence images, in the presence of 0.1 μM PD0325901 or U0126, as compared to DMSO. **(e)** Representative Oct4:GFP FACS for cells in d. **(f)** Oct4:GFP MFI for cells in d. **g**. Model depicting the narrow ERK activity range that limits the number of reprogramming cells to be low, represented by activated Oct4:GFP.

