## supplemental methods for "Morphomechanic tuning of ERK by actin-TFII-IΔ regulates cell identity"

### Supplemental Materials (with full description of materials and procedures)

Four supplementary tables (Table S1-4) and movies (Movie S1-4) accompany this manuscript. Supplementary Table S5 contains sequences for all primers used.

Supplementary materials and methods.

### Mice and Cells

All mouse work was approved by the Institutional Animal Care and Use Committee of Yale University. The reprogrammable mice with reporter (R26rtTA;Col1a14F2A;Oct4GFP) were derived by crossing reprogrammable mice with Oct4:GFP mice, which has been described before.[1-3] Mouse Embryonic Fibroblasts (MEFs) were derived from E13.5 embryos. Feeder cells were obtained by irradiating P5-P6 MEFs. 293T, MEFs and human secondary fibroblasts[4,5] were cultured in DMEM (Gibco, 11995) supplemented with 10% heat inactivated FBS, Pen/Strep/Glutamine (PSG ,100X). Red blood cell depleted hematopoietic progenitors were isolated from spleen of 3-4 weeks old reprogrammable mice. iPSCs were maintained on feeders with ESC medium: DMEM with 15% of ES grade FBS, non-essential amino-acids, Pen/Strep/Glutamine, 0.1 mM 2-mercaptoethanol and LIF (10e7 U/mL, Millipore, 10000X). ESCs were derived from E3.5 embryos.

### Plasmid Construction, Preparation and Virus Production

NLS-actin was generated by fusing the coding sequence for a FLAG tag (ATGGACTACAAGGACGACGATGACAAG) and a nuclear localization signal (NLS, ATGCCAAAAAAGAAGAGAAAGGTACCTCGAGCT) at the N-terminus of *Actb* (Actin Beta, NM_007393.5, subcloned into the pMSCV-IRES-mCherry FP backbone (addgene 52114). EV controls only have FLAG. pMSCV-IRES-blasticidin was generated by replacing the mCherry with blasticidin from pMSCV-IRES-mCherry FP backbone, All actin constructs were also subcloned into pMSCV-IRES-blasticidin backbone. Blasticidin selectable lentiviral vectors for the Dox-inducible expression of control (WT H3.3) or dominant negative Arp2 NLS-mCherry [6] (dnArp2-NLS-mCherry) vector were described before. LentiGuide-Hygro-mTagBFP2 (addgene #99374) was used for custom CRISPR-cas9 gRNA library generation. The shRNAs targeting *Gtf2i* and *Gtf2iβ* were generated by inserting the short hairpin sequence into the lentiviral backbone psi-LVRU6MP (GeneCopia). All sequences are listed in Supplementary Table 5. The pSFG-GFP, pSFG-TFII-I-GFP delta and pSFG-TFII-I-GFP Y248&249F were purchased from Addgene (#22199, #22190, #22196)[7]. mCherry-fusion TFII-I constructs are in pMSCV backbone. Recombinant GST-TFII-I-2D9B, GST-TFII-I-2DN4, GST-TFII-I-2ED2 and GST-TFII-I-2EJE were generated in the pGEX6P vector using the pSFG-TFII-I-GFP delta as the PCR templates. All plasmid DNA was sequence verified. Retrovirus and lentivirus were prepared by transfecting plasmids following maxi or midi preparations (Qiagen, #28704, #12943) into 293T cells, using the FuGene™ 6 transfection reagent (Roche), following manufacturer protocol.

### Reprogramming and Analyses

For reprogrammable MEFs, reprogramming was induced by adding Dox to the ESC medium at a final concentration of 2 µg/mL. Viruses were generally transduced or co-transduced one day before changing ESC medium with Dox. Pluripotent colony scoring was described as previously [8]. For colony counting experiments, cells were seeded on feeders in ESC medium and Dox on reprogramming day4. Alkaline phosphatase (AP) staining was performed with an Alkaline Phosphatase Staining Kit II (Stemgent, Cat.#00-0055) according to manufacturer's recommendations. Colony scoring was done on reprogramming day 10.

For Oct4:GFP analysis by FACS, RNA-Seq and Co-IP experiments, reprogramming was performed without feeders. For inhibitor treatments, PD0325901 (Stemcell, 72184) and U0126 (Stemcell, 73522) cells were induced for 3 days before different concentration of inhibitors were added.

### DAPI staining and Immunofluorescence

To measure nuclear height or colony height, reprogrammable MEFs were cultured in µ-Slide 8 well chambers (ibidi GmbH, Martinsried, Germany), washed twice with PBS, fixed with 4% paraformaldehyde (PFA) for 10 minutes. DNA was labeled by DAPI staining. Z-stack images were taken on a Leica SP5 or Stellaris confocal microscope (Leica Microsystems, Germany). ImageXpress Micro 4 Imaging system was used for brightfield and Oct4:GFP imaging.

For immunofluorescence, cells were fixed, washed in PBS and blocked with 1% BSA, 5% goat serum in PBST for 1 hour at room temperature (RT), and incubated in the diluted primary antibodies overnight at 4°C. Cells were washed twice with PBS and incubated with the appropriate secondary antibodies for 1 hour at RT. DNA was labeled by DAPI staining. For nuclear to cytoplasm ratio measurement, nuclear polylines were drawn along the edge of the nuclei based on DAPI label. The mean pixel values of nuclear fluorescence of Alexa Fluor 488 labeled FLAG antibody were determined using LAS AF software (Leica Microsystems, Germany). The mean pixel values of cytoplastic fluorescence were determined by the area outside of the DAPI labeled nucleus. To measure colocalization of Alexa Fluor 488 labeled FLAG and Alexa Fluor 647 labeled TFII-I, JACoP plug-in[9,10] on Fiji software was used.

### Preparation Of Cell Lysates and Western Blot Analysis

For whole cell lysates, cells were harvested by directly lysing the cells with 2x Laemmli sample buffer (bio-red, P1610747). To prepare cytoplasmic lysate (C) and nuclear lysate (N), identical numbers of cells were harvested. For cytoplasmic protein extraction, hypotonic lysis buffer (10 mM HEPES, with 1.5 mM MgCl2 and 10 mM KCl, freshly added 1mM DTT and protease inhibitors) was used to swell the cell. After 15 minutes incubation on ice, 0.6% IGEPAL CA-630 was added, centrifuged immediately for 30 seconds at 10,000-11,000 x g. The supernatant is cytoplasmic fraction. Nuclear extraction buffer (20 mM HEPES, with 1.5 mM MgCl2, 0.42 M NaCl, 0.2 mM EDTA, and 25% (v/v) Glycerol, freshly added 1mM DTT and protease inhibitors) was added to the pellet, incubated on ice for 30 minutes with gentle shake, centrifuged for 5 minutes at 20,000-21,000 x g. The supernatant is nuclear fraction. All protein samples were denatured in Laemmli buffer and heated at 95°C for 5 min before loading to immunoblot analysis.

Proteins were separated by Mini-Protean TGX Stain Free Gels 4-20% (Bio-Rad, 4568096) alongside Precision Plus Protein Dual Color Standards (Bio-Rad, 1610374) by SDS-PAGE. Once separated, the proteins were transferred onto a PVDF membrane (Bio-Rad) using Bio-Rad Trans-Blot Turbo for immunoblotting. The membranes were blocked with 5% nonfat dry milk in TBS-Tween (TBST) for 1 hour, incubated with primary antibodies overnight at 4  C, followed by incubation with horseradish-peroxidase-conjugated secondary antibodies for 1 hour, and illuminated by enhanced chemiluminescence (ECL). The primary antibodies information is provided in Supplementary Table 5. ECL signal was visualized with enhanced chemiluminescence and measured. Semi-quantification of band intensity was done in Fiji software. The grayscale intensity of each band in the images of the protein blot was measured, the nuclear enrichment was calculated using nuclear to cytoplasmic ratio.

**FACS Analysis and Sorting**

Cells were dissociated with 0.25% Trypsin-EDTA, washed and resuspended with DPBS before FACS analysis or sorting. CD45-APC conjugated antibody (BD Biosciences, San Jose, CA, #559864) was used to label the hematopoietic cells. Cells were analyzed on BD LSRII or sorted on BD FACS Aria Cell Sorter. FlowJo software was used for all FACS plotting and measure the mean fluorescence intensities (MFI) of the Oct4:GFP positive population.

**Co-immunoprecipitations (Co-IP) with MS/MS analysis**

For the reprogrammable MEFs transduced by EV, WT actin or NLS-actin constructs, all mCherry positive populations were sorted by Aria cell sorter. Cells were cultured and collected 6 days after dox induction. For the nuclear input preparation, cytoplasmic fractions were isolated and removed as described in preparation of cell lysates. The nuclear pellets were washed once by micrococcal nuclease buffer (10 mM Tris-HCl, 1 mM CaCl2). For nuclear pellets from 4 $x10^8$ cells, 5 µl micrococcal nuclease (NEB # M0247S) was used, incubated for 15 min at 37°C. Digestion was stopped by adding 10 µl of 0.5 M EDTA. The digested nuclei were pelleted by centrifugation at 10,000-11,000xg for 1 min at 4°C. Nuclear extraction buffer was added to the pellet, incubated on ice for 30 minutes with gentle shake. The lysate was sonicated with 3 sets of 1min pulses at 30 sec intervals, centrifuged for 5 minutes at 20,000-21,000 x g. Balance buffer (20 mM HEPES, with 1.5 mM MgCl2, 0.2 mM EDTA, and 25% (v/v) Glycerol, freshly added 1mM DTT and protease inhibitors) was added to adjust the NaCl to final 150mM. Supernatant/digested protein preparation was transferred to a new tube as the nuclear input for co-immunoprecipitation. For co-immunoprecipitation, the micrococcal nuclease digested protein preparation was diluted 5X by low salt wash buffer (150mM NaCl, 50 mM Tris-HCl), incubate with anti-FLAG m2 magnetic beads (sigma, M8823) for overnight at 37°C with rotation. The beads were washed for 3 times with low salt wash buffer, once with high salt wash buffer (500mM NaCl, 50 mM Tris-HCl), once with pre-urea wash buffer (50 mM Tris, 1 mM EGTA, 75 mM KCl) at 4°C for 5 min with rotation. Elution of the pull-down proteins was by 200 ul urea elution buffer (6–8 M Urea, 20 mM Tris pH 7.5, and 100 mM NaCl) and rotate for 30 min at room temperature. Repeat elution once and pool the eluate. A small portion of unconcentrated proteins was set aside for western blotting and Silver Stain (Pierce, #24600). Most of eluate protein was concentrated by Amicon Ultra-0.5 Centrifugal Filter Devices (Millipore, Z677906). The unconcentrated and concentrated protein elations were denatured in Laemmli buffer and heated at 95°C for 5 min before loading to gels. For mass spectrometry (MS), gels were stain by GelCode® Blue Stain Reagent (Thermo Scientific 24590) and the whole lanes were cut and analyzed by the Keck Proteomics Center at Yale to identify the pull-down proteins. STRING analyses were applied for explore the function of potentially interacted proteins.

**Custom Library Generation and CRISPR-Cas9 Screen**

The online web tool CHOPCHOP (https://chopchop.cbu.uib.no/) was used to generate sgRNA designs against target genes[11]. For each gene, 4 sgRNAs were chosen based on the location and score. Screening is done following Feng Zhang Lab's protocols with minor modification [12,13]. Briefly, for each sgRNA, add 5'-CACCG and delete 3'-NGG. Reverse complement of the first oligo without 5'-adaptor and 3'-NGG, add 5'-AAAC and 3'-C-3 to reversed sequence as the second oligo. Oligo pairs were ordered from Integrated DNA Technologies (IDT). Each pair of oligos were annealed separately and then pooled together in one tube for phosphorylation and ligation. LentiGuide-Hygro-mTagBFP2 were digest by BsmBI and ligated with the oligo pool, transformed into MAX Efficiency™ DH5α-T1R Competent Cells (Invitrogen, #12034013) following the manufacturer protocol. Transformed bacteria were plated on LB-AMP plates and incubated at 37°C overnight. All bacteria colonies were counted and harvested with LB medium. At 22-fold coverage, over 12,000 bacteria colonies were mixed for plasmid preparation. The plasmid DNA was isolated (Qiagen, #12943) following the manufacture's protocol. Lentivirus were made by general protocols, and 50X concentrated lentivirus was used for screen. For screen, reprogrammable MEFs transduced by three constructs lentiCas9-Blast (Addgene, #181025), the gRNA library and NLS-actin (co-expressing mCherry). Input DNA was collected 48h after transduction of custom gRNA library. After selection by blasticidin, all the Cas9 positive cells were sorted. The BFP and mCherry double positive populations and BFP only populations were separated for culture. DNA was collected from the reprogramming day4 cells after dox induction. The input and day4 cells all has >35x coverage per sgRNAs. All DNA was isolated (Qiagen, #51104) following the manufacture's protocol. PCR by choice taq blue mastermix (Denville, CB4065-7) using the barcoding sequencing primer pairs (Table S4). PCR products were run on 2% agarose gel, and size was selected by gel electrophoresis and purified (Qiagen, #28704). Quality control of DNA was done by Qubit fluorometer (Invitrogen) and Agilent 2100 Bioanalyzer. Samples that passed quality control were used for sequencing with the Illumina NovaSeq System. For data processing for CRISPR screens, reads that pass QC were preprocessed by locating the first 8 bp barcode of one of the three anchors used in the barcoding primers and extracting the 20 bp preceding the anchor using a bespoke Perl script. Most of the 20 bp trimmed reads matched the designed library. Read counts from input, Double and BFP in the screen were combined in a matrix. Data analysis was performed with the MAGeCK, MAGeCK-VISPR and MAGeCKFlute software[14-16]. In brief, MAGeCK in Python was used to normalize the screen from input. MAGeCK-VISPR calculated the 'beta score' for each gene. MAGeCKFlute pipeline in R, which was designed to perform quality control, normalization, and downstream analysis of the functional CRISPR screens, was used to identify differentially enriched genes in the presence or absence of nuclear actin. Loess (loess Boolean, specify whether to include loess normalization in the pipeline) method was chosen for the normalization.

**GST Pull-down Assay**

The plasmids for GST-TFII-I-2D9B, GST-TFII-I-2DN4, GST-TFII-I-2ED2 and GST-TFII-I-2EJE were transformed into E. coli BL21 gold (DE3). 50ml of E. coli were induced by 0.55 mM IPTG for 7 hours at room temperature, lysis buffer (50 mM Tris-HC pH 7.5, 300 mM NaCl, 10% Glycerol, freshly add 5mM 2-mercaptoethanol and protease inhibitor cocktail) were added and sonication were performed to lysis the bacteria. The GST-fusion proteins in supernatant of bacterial lysate were incubated with glutathione agarose beads for 2 hours at 4°C under gentle rotation. After 3 washes with PBS, EV or NLS-Actin transduced 293T cell lysates in RIPA buffer were added to GST-fusion proteins immobilized on the beads. The EV/NLS-actin lysates and the immobilized GST-fusion protein bound beads were incubated overnight at 4°C under gentle rotation. After three washes, the proteins bound to beads were eluted and analyzed by immunoblotting. For GST pull-down, GST-2EJE or GST control were immobilized to glutathione agarose beads. Immobilized beads were used to incubate with or without G-actin for 3h. G-actin was prepared from rabbit skeletal muscles as describe before [17,18]. The GST-pulldown and flow-through supernatant were analysis by SDS-PAGE and GelCode® Blue Stain Reagent.

**Semi-quantitative PCR and Quantitative PCR**

Total RNA was extracted with Trizol® reagent (Invitrogen) and reverse transcribed with the SuperScript III First-Strand Synthesis System (Invitrogen) according to manufacturer instruction. PCR reactions were set up in triplicates with iQ™ SYBR® Green Supermix (Biored, # 1708880) with 28 cycles. PCR products were run on 2% agarose gel. Quantification of band intensity was done in ImageJ software. The grayscale intensity of each band was measured, the Relative *Gtf2iΔ* expression was calculated by calculating the ratio of smaller band to bigger band ratio. Sanger sequencing was performed to validate the smaller band to be *Gtf2iΔ*, and the larger band to be *Gtf2iβ*. Real-time qPCR was performed on a Bio-Rad CFX96 or CFX384.

**Live-Cell Imaging and ERK Activity Assay**

For live-cell imaging, ImageXpress Micro 4 Imaging system was used with environmental control at 37°C, compressed gas of 5% $CO_2$, 21% $O_2$ and balance Nitrogen. Time-lapse fluorescence imaging of ERK-KTR-Clover signal were collected over 80 min at 10-minute intervals. For quantitation analysis, Fiji software was used to measure the mean intensity of nucleus(N) and cytoplasm(C) in each time point. The logarithmic ratio of cytoplasmic to nuclear fluorescence intensity, adjusted for background fluorescence: $LOG((C - background)/(N - background))$ was computed for each timepoint, and served as a proxy for ERK. For normalize to time 0, ERK activity at time 0 was subtracted from the ERK activity of all later time points.

**Fluorescence Recovery After Photobleaching (FRAP) Assay**

Reprogrammable MEFs were co-transduced with EV or NLS-actin with TFII-I-Δ-mCherry fusion protein. Double positive cells were sorted and seeded in μ-Slide 8 well chambers. FRAP assay was performed using Leica Stellaris confocal microscope. All experiments were conducted at 37 °C with 5% $CO_2$. Pre-bleach images were acquired by averaging three consecutive images.

2 µm × 2 µm region of interest (ROI1) was gated for bleaching thee mCherry signal on the nuclei, with the WLL laser at 100% intensity. ROI2 signal on the whole fluorescent nuclei and ROI3 signal of the background were collected for normalization. For image collection, mCherry signal of all ROIs at 2% intensity was used, 10 times of pre-bleach, 50 times of bleach and 40 times of post-bleach images were collected at 0.189 s intervals. The easyFRAP application was used for data normalization and processing.[19]

### Statistical and Graphing

All statistical analyses were conducted using Graphpad Prism9, employing t-test or ANOVA based on the comparison group numbers. All graphs were generated using Graphpad Prism9.


1 Eminli, S. *et al.* Differentiation stage determines potential of hematopoietic cells for reprogramming into induced pluripotent stem cells. *Nature Genetics* **41**, 968-U929 (2009). https://doi.org/10.1038/ng.428

2 Megyola, C. M. *et al.* Dynamic migration and cell-cell interactions of early reprogramming revealed by high-resolution time-lapse imaging. *Stem Cells* **31**, 895-905 (2013). https://doi.org/10.1002/stem.1323

3 Carey, B. W., Markoulaki, S., Beard, C., Hanna, J. & Jaenisch, R. Single-gene transgenic mouse strains for reprogramming adult somatic cells. *Nat Methods* **7**, 56-59 (2010). https://doi.org/10.1038/nmeth.1410

4 Hockemeyer, D. *et al.* A drug-inducible system for direct reprogramming of human somatic cells to pluripotency. *Cell stem cell* **3**, 346-353 (2008). https://doi.org/10.1016/j.stem.2008.08.014

5 Maherali, N. *et al.* A high-efficiency system for the generation and study of human induced pluripotent stem cells. *Cell stem cell* **3**, 340-345 (2008). https://doi.org/10.1016/j.stem.2008.08.003

6 Tsopoulidis, N. *et al.* T cell receptor-triggered nuclear actin network formation drives CD4(+) T cell effector functions. *Sci Immunol* **4** (2019). https://doi.org/10.1126/sciimmunol.aav1987

7 Hakre, S. *et al.* Opposing functions of TFII-I spliced isoforms in growth factor-induced gene expression. *Mol Cell* **24**, 301-308 (2006). https://doi.org/10.1016/j.molcel.2006.09.005

8 Hu, X. *et al.* MKL1-actin pathway restricts chromatin accessibility and prevents mature pluripotency activation. *Nat Commun* **10**, 1695 (2019). https://doi.org/10.1038/s41467-019-09636-6

9 Cordelieres, F. P. & Bolte, S. in *ImageJ User & Developer Conference.* 181.

10 Bolte, S. & Cordelières, F. P. A guided tour into subcellular colocalization analysis in light microscopy. *Journal of microscopy* **224**, 213-232 (2006).

11 Labun, K. *et al.* CHOPCHOP v3: expanding the CRISPR web toolbox beyond genome editing. *Nucleic acids research* **47**, W171-W174 (2019). https://doi.org/10.1093/nar/gkz365

12 Sanjana, N. E., Shalem, O. & Zhang, F. Improved vectors and genome-wide libraries for CRISPR screening. *Nat Methods* **11**, 783-784 (2014). https://doi.org/10.1038/nmeth.3047

13 Shalem, O. *et al.* Genome-scale CRISPR-Cas9 knockout screening in human cells. *Science* **343**, 84-87 (2014). https://doi.org/10.1126/science.1247005

14 Li, W. *et al.* MAGeCK enables robust identification of essential genes from genome-scale CRISPR/Cas9 knockout screens. *Genome Biol* **15**, 554 (2014). https://doi.org/10.1186/s13059-014-0554-4
15 Wang, B. *et al.* Integrative analysis of pooled CRISPR genetic screens using MAGeCKFlute. *Nature Protocols* **14**, 756-780 (2019). https://doi.org/10.1038/s41596-018-0113-7
16 Li, W. *et al.* Quality control, modeling, and visualization of CRISPR screens with MAGeCK-VISPR. *Genome Biol* **16**, 281 (2015). https://doi.org/10.1186/s13059-015-0843-6
17 Cao, W., Goodarzi, J. P. & De La Cruz, E. M. Energetics and kinetics of cooperative cofilin-actin filament interactions. *J Mol Biol* **361**, 257-267 (2006). https://doi.org/10.1016/j.jmb.2006.06.019
18 Kang, H. *et al.* Site-specific cation release drives actin filament severing by vertebrate cofilin. *Proc Natl Acad Sci U S A* **111**, 17821-17826 (2014). https://doi.org/10.1073/pnas.1413397111
19 Giakoumakis, N. N., Rapsomaniki, M. A. & Lygerou, Z. Analysis of Protein Kinetics Using Fluorescence Recovery After Photobleaching (FRAP). *Methods Mol Biol* **1563**, 243-267 (2017). https://doi.org/10.1007/978-1-4939-6810-7_16
